# In defense of Apocynaceae: inference on evolution of pyrrolizidine alkaloids from evolution of an enzyme in their biosynthetic pathway, homospermidine synthase

**DOI:** 10.1101/2023.04.01.535177

**Authors:** Chelsea R. Smith, Elisabeth Kaltenegger, Jordan Teisher, Abigail J. Moore, Shannon C. K. Straub, Tatyana Livshultz

## Abstract

**Premise:** When enzymes encoded by paralogous genes produce identical specialized metabolites in distantly related plant lineages, it is strong evidence of parallel phenotypic evolution. Inference of phenotypic homology for metabolites produced by orthologous genes is not so straightforward, however, since orthologs may be recruited in parallel into novel pathways. Prior research on pyrrolizidine alkaloids (PAs), specialized metabolites of Apocynaceae, reconstructed evolution of homospermidine synthase (HSS), an enzyme of PA biosynthesis, and inferred a single origin of PAs because HSS enzymes of all known PA-producing Apocynaceae species are orthologous and descended from an ancestral enzyme with the predicted motif (VXXXD) of an optimized HSS.

**Methods:** We increased sampling, tested the effect of amino acid motif on HSS function, revisited motif evolution, and tested for selection to infer evolution of HSS function and its correlation with phenotype.

**Key results:** Some evidence supports a single origin of PAs: an IXXXD HSS, similar in function to VXXXD HSS, evolved in the shared ancestor of all PA-producing species; loss of optimized HSS occurred multiple times via pseudogenization and perhaps via evolution of an IXXXN motif. Other evidence indicates multiple origins: the VXXXD motif, highly correlated with the PA phenotype, evolved two or four times independently; the ancestral IXXXD gene was not under positive selection while some VXXXD genes were; substitutions at sites experiencing positive selection occurred on multiple branches in the *HSS*-like gene tree.

**Conclusions:** Complexity of the genotype-function-phenotype map confounds inference of PA homology from *HSS* evolution in Apocynaceae.

---

Identification of the enzymes that synthesize plant specialized metabolites (small molecules that mediate ecological interactions) and reconstruction of their evolution have provided key insights into the origins of phenotypic novelty and convergence. Many enzymes of plant specialized metabolism have been recruited from conserved genes of primary metabolism via gene duplication followed by neo- or subfunctionalization (Weng, 2014; Moghe and Last, 2015; Lou et al., 2022). In several notable cases, e.g. caffeine (Huang et al., 2016) and betalains (Sheehan et al., 2020), identical specialized metabolites in related plant lineages are synthesized by enzymes encoded by paralogs recruited from the same gene families, for more examples see (Pichersky and Lewinsohn, 2011), thus explaining the pattern of high but imperfect correlation between plant phylogeny and plant specialized metabolism frequently noted by chemotaxonomists (Kubitzki and Gottlieb, 1984; Wink, 2003). While paralogy of genes involved in synthesis of identical specialized metabolites shared by distantly related taxa has been interpreted as strong evidence of convergent phenotypes (Pichersky and Lewinsohn, 2011; Lou et al., 2022), the corollary: orthology of genes involved in synthesis of identical specialized metabolites shared by (relatively) closely related taxa as evidence of homologous phenotypes, has not been considered as carefully.

There is little ambiguity about homology of specialized metabolites when they are synthesized by sister taxa with enzymes encoded by orthologous genes. But when there is a high level of homoplasy in the distribution of a metabolite among closely related taxa, combined with the understanding that duplicated genes may persist for millions of years before being lost or evolving new functions (Panchy et al., 2016), the possibility must be considered that orthologous copies of a gene were independently recruited into convergently evolved metabolic pathways. The balance of evidence tips toward homology of phenotypes if it can be shown that 1) there is tight correlation among gene sequence, enzyme function, and phenotype, 2) the novel function of the enzyme recruited to the novel metabolic pathway evolved once in the ancestral gene, and 3) patterns of selection on the gene are consistent with emergence of a novel adaptive phenotype (i.e. positive selection) in the common ancestor of all species that have the phenotype today followed by its loss in the species that do not (i.e. neutral evolution).

Pyrrolizidine alkaloids (PAs), often toxic (Stegelmeier et al., 1999; Tamariz et al., 2018) specialized metabolites, are characterized by a necine base, derived from homospermidine, with one or more attached necic acids (Hartmann and Witte, 1995; Ober and Hartmann, 2000). PAs likely evolved as a defense against insect herbivores (Hartmann, 1999; Ober and Kaltenegger, 2009) and occur in 15 angiosperm families (Appendix S1,Table S1; see Supplemental Data with this article) (Hartmann and Witte, 1995; Nawaz et al., 2000; Kim et al., 2010; Tamariz et al., 2018; Barny et al., 2021), including Apocynaceae where they are present in three or four of the sub-lineages (Nerieae, Echiteae, Apocyneae, and potentially Malouetieae) within a well-supported lineage (the APSA clade) (Fig. 1) (Barny et al., 2021).

**Fig. 1:**
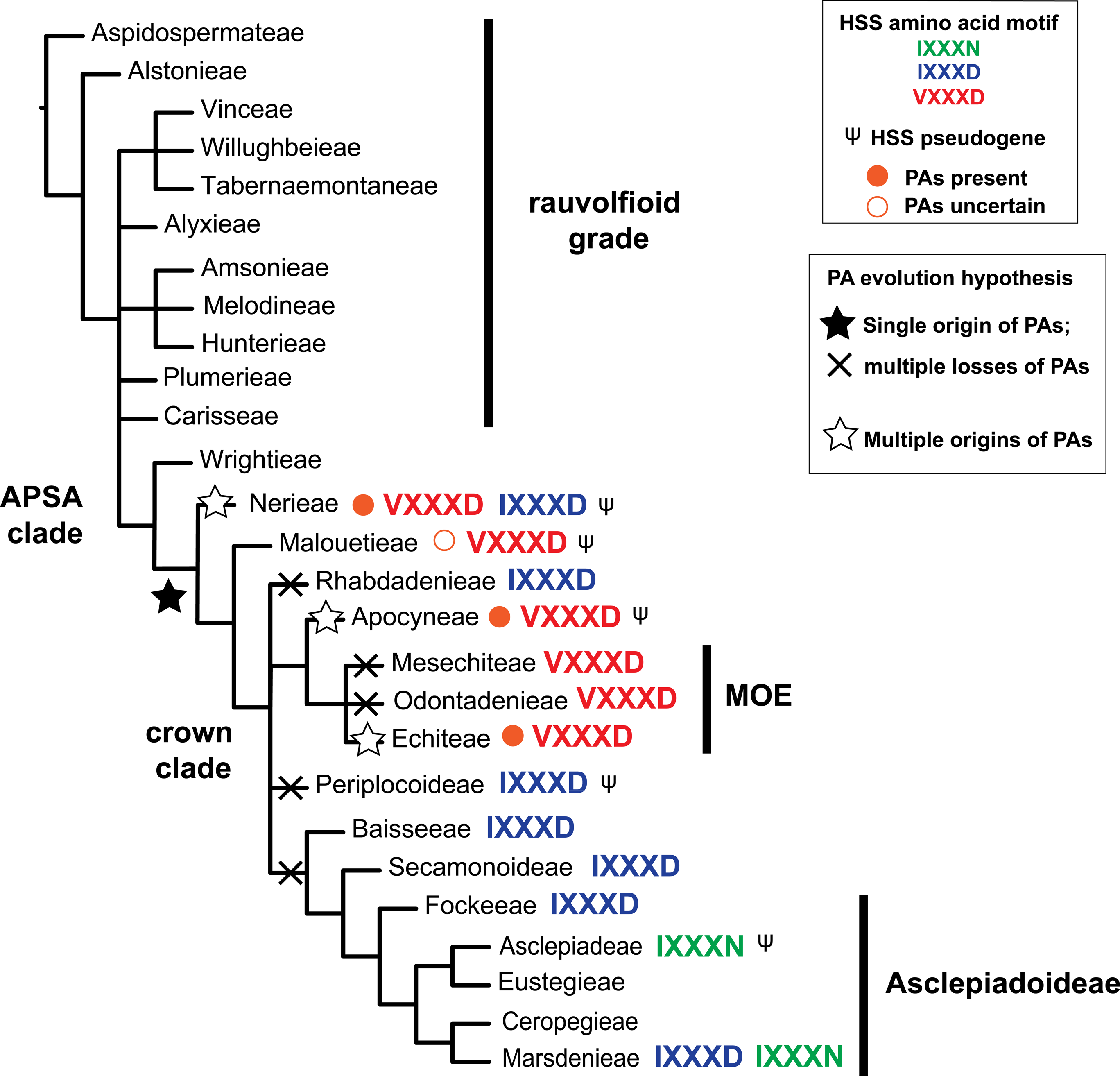
Phylogenetic relationships among Apocynaceae subfamilies and tribes, with pyrrolizidine alkaloid (PA) distribution (orange circles), and distribution of motifs and pseudogenes (Ψ) among *HSS*-like genes mapped. Topology reflects statistically significant relationships supported by the polytomy test in the ASTRAL analysis of Antonelli et al. (2021) based on nuclear gene trees, with addition of Odontadenieae, placed here in a polytomy with Mesechiteae and Echiteae (Livshultz et al., 2007; Livshultz, 2010; Morales et al., 2017; Fishbein et al., 2018) and Carisseae, placed here in the rauvolfioid polytomy (Fishbein et al., 2018). Orange circles at tips represent the presence of PAs with high confidence as defined in Barny, et al. (2021) in at least one species of the group. The amino acid motif (positions 269 and 273) of *HSS*-like genes is indicated at tips (VXXXD in red, IXXXD in blue, and IXXXN in green). Nerieae has multiple motifs (VXXXD in *Strophanthus* and *Isonema*, IXXXD in *Alafia*; Marsdenieae has two *HSS*-like paralogs with IXXXD and IXXXN motifs (see Fig. 3). Ψ indicates that at least one species in the lineage has only a pseudogenized copy of *HSS*. Alternative PA evolution hypotheses are mapped on the branches: black star represents the single origin of PAs in the MRCA of PA-producing lineages followed by multiple losses (black X’s) in lineages not currently known to produce PAs. Alternatively, the white stars represent multiple origins of PA biosynthesis in the MRCA of each PA-producing lineage.

Clarifying whether PAs are homologous or convergent among Apocynaceae lineages is important for chemotaxonomy of the family: are they a synapomorphy for the APSA clade even if rare and scattered within it (Fig. 1)? And also to test a co-evolutionary hypothesis for the origin of PA-pharmacophagy [adult acquisition of chemicals from non-food sources (Boppré, 1984)] in the PA- and Apocynaceae-philous milkweed and clearwing butterflies (Lepidoptera subfamily Danainae) which includes the North American monarch (*Danaus plexippus*) and queen (*D. gillipus)* butterflies (Brower et al., 2010; Brower et al., 2014). Danainae are used as a model system to study adaptation of specialist herbivores to host plant chemistry (Petschenka and Agrawal, 2015, 2016). Edgar (1984) proposed a “defense de-escalation” hypothesis (Livshultz et al., 2018): that the most recent common ancestor (MRCA) of Danainae first acquired PAs via larval sequestration from a PA-producing Apocynaceae species, some of whose descendants, under selection from their PA-adapted herbivores, lost PA-production, causing Danainae to continue to feed on their now PA-free host plants as larvae but to obtain PAs from other sources as adults. Notably, 702 of 726 larval host plant records for Danainae on Apocynaceae are from species in the APSA clade (Livshultz et al., 2018). The MRCA of these larval host plant species is also the MRCA of all known PA-producing Apocynaceae species (Livshultz et al., 2018) (Fig. 1).

Because distribution of PAs in Apocynaceae is underdetermined (Burzynski et al., 2015; Barny et al., 2021, 2022), Livshultz et al. (2018) studied a gene in their biosynthetic pathway to test the defense de-escalation hypothesis. Biosynthesis of PAs begins with homospermidine synthase (HSS). Paralogy of *HSS* genes in distantly related plant families has been interpreted as strong evidence that PA biosynthesis evolved repeatedly among angiosperms (Reimann et al., 2004). HSS evolved via gene duplication from deoxyhypusine synthase (DHS) (Ober and Hartmann, 2000), a highly conserved enzyme of eukaryotic primary metabolism that activates eIF5A, a translation elongation factor (Chen and Liu, 1997; Park et al., 2010) (Fig. 2a), by transferring an aminobutyl group from spermidine to a lysine residue in eIF5A to form deoxyhypusine. HSS has lost this function but has retained and optimized another function of DHS: catalysis of putrescine with spermidine, producing an order of magnitude more homospermidine than DHS (Fig. 2b) (Ober et al., 2003b; Reimann et al., 2004; Kaltenegger et al., 2013). In addition to catalyzing the reaction of putrescine with spermidine to produce homospermidine and 1,2 diaminopropane (Fig. 2c, reaction I), HSS can also catalyze the reaction of two spermidine molecules to produce canavalmine and 1,2 diaminopropane (Fig. 2c, reaction II), and the lysis of spermidine to produce 1,2 diaminopropane and the Δ^1^-pyroline (Fig. 2c, reaction III) (Kaltenegger et al., 2021). Eight independent *DHS/HSS* duplications occurred in seven PA-producing angiosperm families (Anke et al., 2004; Reimann et al., 2004; Kaltenegger et al., 2013; Gill et al., 2018; Livshultz et al., 2018), consistent with at least eight independent origins of pyrrolizidine alkaloids.

**Fig. 2:**
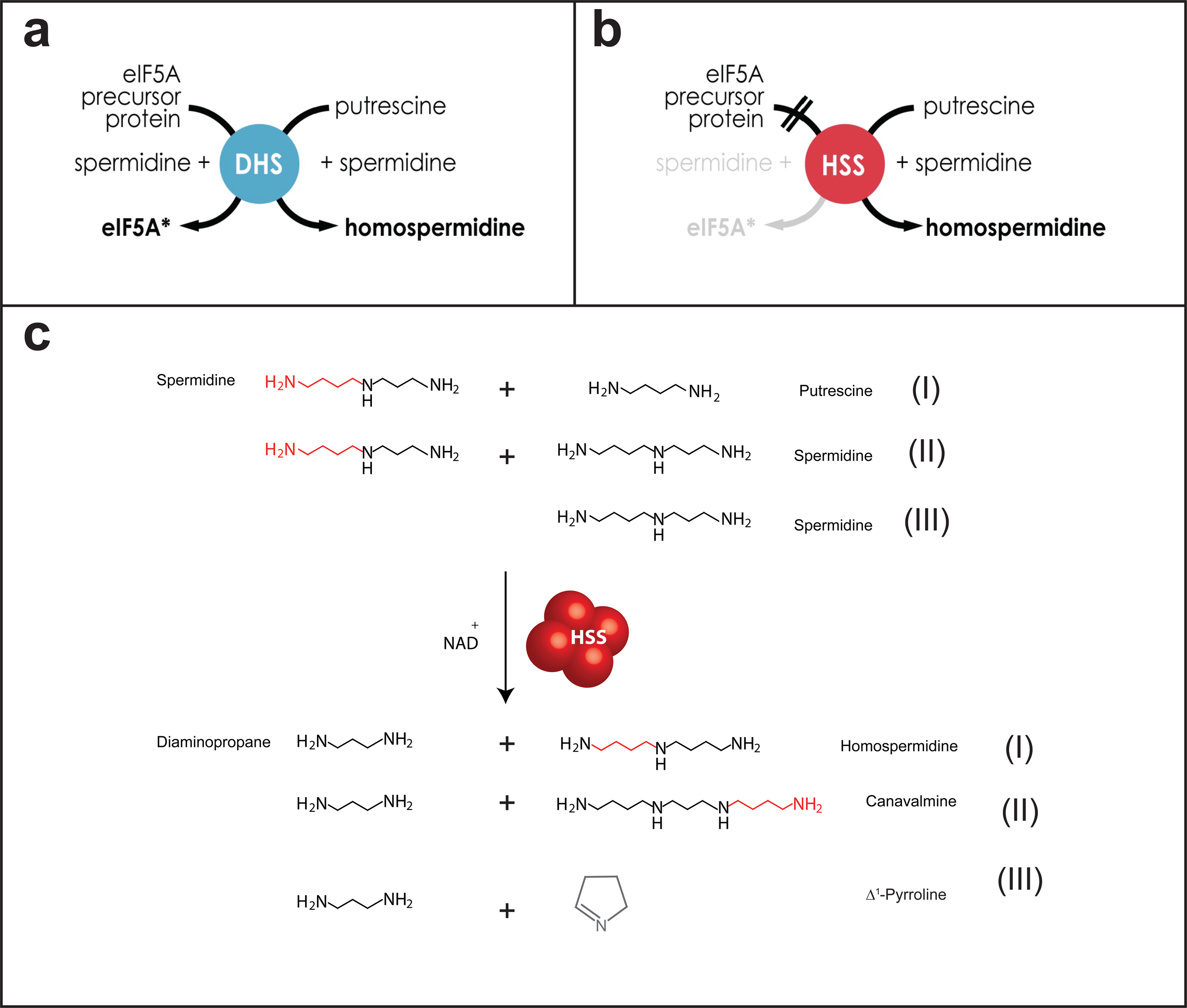
Function of deoxyhypusine synthase (DHS, a) and homospermidine synthase (HSS, b, c). **a.** DHS activates the eIF5A translation elongation factor by transferring the amino-butyl moiety of spermidine to a lysine residue on the eIF5A precursor protein to form deoxyhypusine (Chen and Liu, 1997). DHS can also transfer the amino-butyl moiety of spermidine to putrescine to form homospermidine (Ober and Hartmann, 1999). **b.** HSS (homospermidine synthase) has lost activity with eIF5A, but has maintained and optimized activity with putrescine, producing an order of magnitude more homospermidine than DHS (Ober et al., 2003a; Reimann et al., 2004; Kaltenegger et al., 2013). Homospermidine is precursor to the necine base of a pyrrolizidine alkaloid. **c**. Reactions catalyzed by HSS showing substrates and products.

Prior research suggested a tight coupling of *HSS* genotype and PA phenotype. Livshultz, et al. (2018) showed that an HSS-specific amino acid motif (VXXXD) evolved from a DHS-specific motif (IXXXN) in each of the seven independent origins of HSS and PAs among the angiosperm families they analyzed. Further evidence for the functional importance of this HSS motif derives from natural infra-specific polymorphisms (Gill et al., 2018; Prakashrao et al., 2022) and experimental mutagenesis (Kaltenegger et al., 2013; Prakashrao et al., 2022). Evidence that the motif was under positive selection in the evolution of HSS from DHS in Convolvulaceae also supported its functional importance and correlation with phenotypic evolution (Kaltenegger et al., 2013).

Livshultz et al. (2018) showed that the pattern of *DHS/HSS* duplication, IXXXN/VXXXD motif evolution, and pseudogenization in Apocynaceae supports predictions of the defense de-escalation hypothesis: that PA biosynthesis evolved once in Apocynaceae and was lost in multiple lineages (Fig. 1). They showed a single *DHS/HSS* duplication in the most recent common ancestor (MRCA) of all PA-producing taxa in Apocynaceae, early in the diversification of the APSA clade, and an ancestral HSS with a VXXXD motif. HSS enzymes in genera known for PAs retain the VXXXD motif with one exception [*Alafia* (Nerieae), IXXXD, although the sequenced species tested negative for PAs, and PA-positive *Alafia* species were not sequenced], while genera not known for PAs either retained the VXXXD motif or evolved IXXXD or IXXXN motifs (Barny et al., 2021, 2022) (Figs. 1, 3). Furthermore, *HSS* is pseudogenized in *Asclepias syriaca* (Asclepiadeae), a well-studied PA-free species (Livshultz et al., 2018) (Fig. 1).

Here, we revisit the evolution of *HSS*-like genes in Apocynaceae with expanded sequencing, functional experiments, phenotypic data (Barny et al., 2021, 2022), and tests for selection to ask if we can substantiate a single ancestral recruitment of HSS to PA biosynthesis followed by multiple independent losses of the PA phenotype (Fig. 1) or if the data better support multiple independent recruitments and multiple independent origins of the PA phenotype (Fig. 1) based on the criteria of 1) correlation of enzyme function, *HSS* genotype, and PA phenotype, 2) phylogenetic placement of the evolution of novel enzyme functions, and 3) selection on enzyme function.

## MATERIALS AND METHODS

### Taxon sampling for DHS/HSS sequencing—

One hundred seventy (170) accessions of 159 Apocynaceae species (Appendix S1, Table S2), including 10 known PA-producing species were sampled (Appendix S1, Table S2). The outgroup was *Gelsemium sempervirens* (Gelsemiaceae) (Antonelli et al., 2021). Previously published Apocynaceae *DHS* and *HSS* sequences from 25 species were also included (Livshultz et al., 2018).

### Calotropis genome query—

*DHS*- and *HSS*-like genes were extracted from the *Calotropis gigantea* genome (Hoopes et al., 2018) via tblastx searches with exon sequences of *Parsonsia alboflavescens DHS* and *HSS* (MG817648.1, MG817649.1) in Geneious Prime v.2020.0.3 (https://www.geneious.com).

### DNA extraction, library preparation, targeted enrichment, sequencing—

DNA extraction, library preparation, targeted enrichment, and paired end sequencing were previously described (Straub et al., 2020). Probes were designed based on the Apocynaceae *DHS* and *HSS* sequences published by Livshultz et al. (2018), manually trimmed to not more than 200 bp beyond the 5’ and 3’ ends of the first and last exons. A total of 2707 probes targeted *DHS* and *HSS* (Straub et al., 2020). Contaminated libraries (i.e. libraries containing DNA from more than one sample) were identified by mapping a sample’s reads to its own assembled plastome. Samples with evidence of two divergent plastomes were considered contaminated and excluded (unpublished data).

### Contig assembly—

Raw paired end sequence reads were trimmed using Trimmomatic default settings (slidingwindow:10:20, minlen:40) (Bolger et al., 2014). Using the first stage of the MyBaits pipeline [BLASTN (Cameron and Williams, 2007), SPAdes (Bankevich et al., 2012)] with default options (SPAdes: k 21,33,55,77, cov-cutoff off, phred-offset 33) (Moore et al., 2018), trimmed sequence reads for each sample were binned as *DHS*/*HSS* by BLASTN using reference transcriptomic *DHS*-like sequences (Appendix S1, Table S2) and binned reads were assembled using SPAdes. Libraries that did not have an assembled contig with > 20x coverage were removed from analysis (Bentley et al., 2008; Dohm et al., 2008; Harismendy et al., 2009; Whittall et al., 2010; Straub et al., 2012). Retained SPAdes contigs were extended and fused by afin, an assembly finishing program (-s 50, −1 100) (McKain and Wilson, 2017).

### Exon annotation—

Retained afin contigs were annotated for exonic regions in Geneious Prime v.2020.0.3 (https://www.geneious.com) using *DHS* and *HSS* exons from *Parsonsia alboflavescens* (GenBank accessions: MG817648.1, MG817649.1) (50% identical threshold). Annotated contigs (exon and intron) were aligned within a sample to identify and manually annotate partial or divergent exons that did not meet identity threshold.

### Alignment construction—

All alignments were constructed as nucleotide alignments using MAFFT v7.450 (gap penalty=3) (Katoh et al., 2002; Katoh and Standley, 2013) in Geneious Prime v.2020.0.3 (https://www.geneious.com).

### Maximum likelihood tree construction—

Maximum likelihood trees were constructed with rapid bootstrapping and partitioned by codon position in RAxML-HPC v.8 on XSEDE (v8.2.12) (GTR+GAMMA, 1000 bootstrap replicates) (Stamatakis, 2014) in CIPRES Science Gateway (v3.3) (Miller et al., 2010).

### Gene assembly from contigs—

Annotated exons were extracted and aligned with an existing Apocynaceae sequence alignment (Livshultz et al., 2018) and *Calotropis gigantea* sequences (Hoopes et al., 2018). A maximum likelihood tree was built from this “initial alignment” (Appendix 2, Appendix 1: Tables S2, S3). Contigs orthologous to *Parsonsia alboflavescens HSS* were considered *HSS*-like, those outside this clade, *DHS*-like. Contigs derived from the same library were then concatenated using the following algorithm. After excluding contigs with non-terminal stop codons, a strict consensus was calculated from overlapping partial contigs that were in a clade with other *DHS*-like or *HSS*-like sequences from the same tribe in the initial alignment gene tree (Appendix S2) from the same library. Amino acid similarity in the region of overlap of the combined contigs ranged from 89.6-100% (Appendix S1, Table S5). Then non-overlapping contigs were concatenated using the same grouping criteria. Polyphyletic contig pairs were grouped if there were no sequences from closely related taxa for them to cluster with, and there was no evidence of contamination in the source library.

### Validation of gene assemblies—

Sequence assemblies for samples that had Sanger sequences of the same locus from Livshultz, et al. (2018) were validated via pairwise alignment and calculation of divergence. Here, and otherwise, divergence/pairwise amino acid sequence identity was calculated in Geneious using in-frame nucleotide alignments; this excludes missing data.

### Construction of matrices—

Sanger sequences >95% identical to new sequences from the same sample were removed from the alignment. The “Full Dataset” consisted of potential pseudogene sequences, full (minimum exons 2-6) *DHS*-like and *HSS*-like sequences, short *DHS*-like and *HSS*-like contigs (from samples in which an entire *DHS*-like or *HSS*-like gene had also been assembled by SPAdes/afin), and non-redundant Sanger sequences from Livshultz, et al. (2018) (Appendix 1, Tables S2, S3). This full dataset alignment was trimmed to produce a “Reduced Dataset” alignment used for ancestral state reconstruction and selection analyses. Potential pseudogenes and sequences that did not span at minimum exons 2-6 were removed from this dataset and remaining sequences were realigned. Thirty-seven (37) bp were trimmed from the 5’ end and 67 base pairs from the 3’ end of this reduced dataset alignment, as these areas were highly variable.

### Gene tree construction—

Three gene trees were produced: a full dataset tree was constructed that contains all contigs (candidate pseudogenes, short contigs, and full/concatenated/consensus contigs) (Appendix S3), and a reduced dataset tree (only containing contigs with at least exons 2-6 and no non-terminal stop codons) (Figs. 3a, b, 4a, Appendix S4). Lastly, a third tree was constructed using the reduced dataset alignment 100% constrained to the full dataset tree topology, referred to as the “full dataset topology” (Figs. 3c, d, 4b, Appendix S5).

**Fig. 3:**
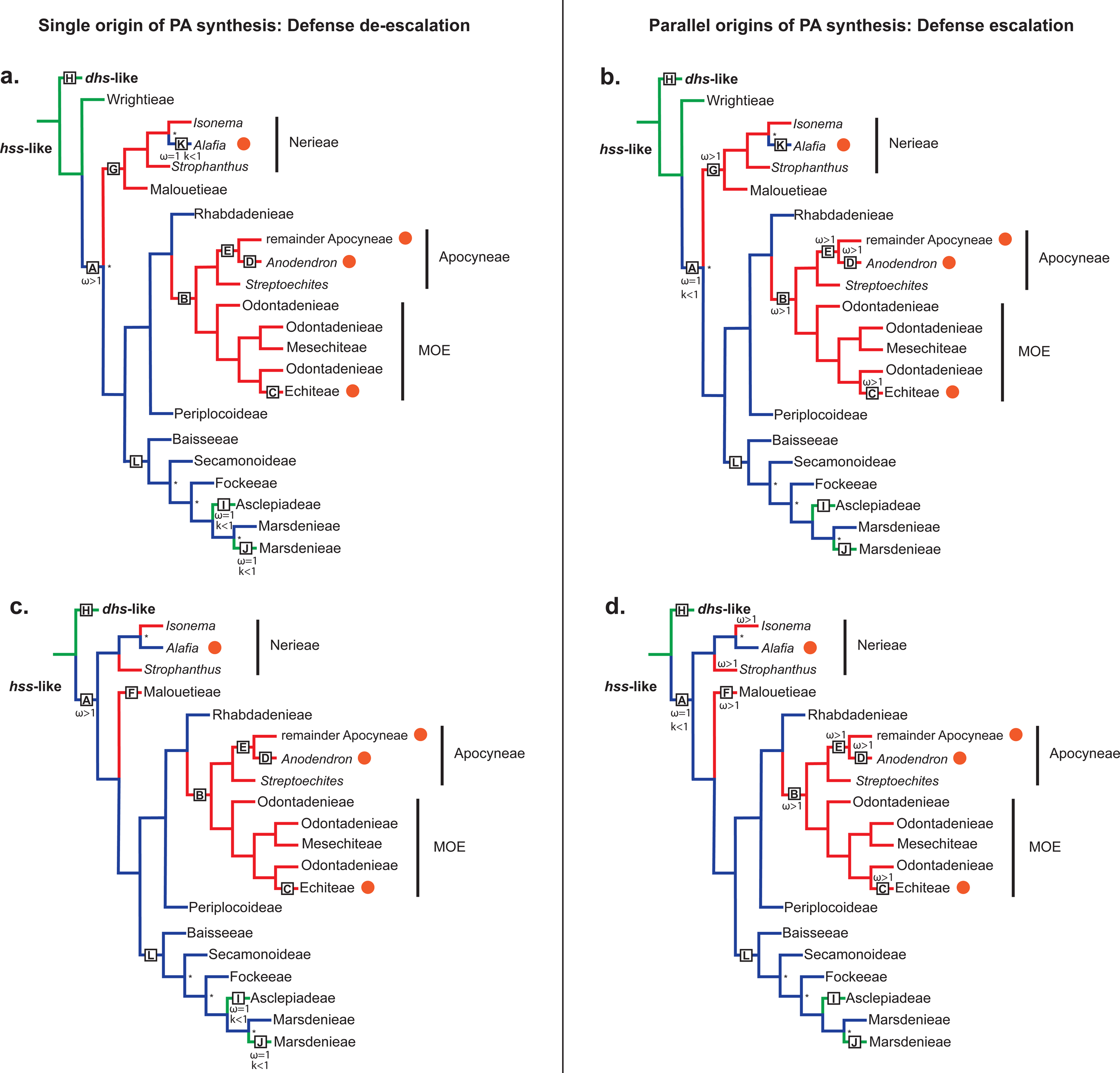
Summary of *DHS/HSS* gene trees based on the reduced dataset topology (subpanels **a, b**, see Appendix S4 for entire tree) and the full dataset topology (subpanels **c, d**, see Appendix S5 for entire tree) with branches colored according to the reconstructed I269VXXXN273D motif (green: IXXXN, blue: IXXXD, red: VXXXD). Distribution of pyrrolizidine alkaloids indicated with orange circles next to the *HSS* gene lineage of the corresponding taxon. The two alternative evolutionary hypotheses of PA biosynthesis and HSS optimization: single origin of PAs followed by multiple independent losses (defense de-escalation, subpanels **a, c**) and multiple parallel origins (defense escalation, subpanels **b, d**) are illustrated on each gene tree topology via the patterns of selection [nonsynonymous/synonymous substitution ratios (ω) and selection intensity (k)] on *HSS* predicted under each scenario. Branches and clades marked by letters (A-L) are used in tests for selection (see Tables 1 and 2). Under the **single origin/defense de-escalation model** (subpanels a, c), PA biosynthesis evolved in the MRCA of all PA-producers, and the ancestral HSS enzyme was optimized under positive selection (Branch A, ω>1). Subsequent parallel losses of PA biosynthesis, under negative or neutral selection, result in neutral evolution of the *HSS* locus (ω=1) and evolution of a more *DHS*-like from a more *HSS*-like motif, i.e. VXXXD to IXXXD in the ancestral *HSS*-like gene of *Alafia* (Branch K, subpanel a) and IXXXD to IXXXN in the ancestral *HSS*-like gene of Asclepiadeae (Branch I, subpanels a and c) and in the IXXXN *HSS*-like paralog of Marsdenieae (Branch J, subpanels a and c). Relaxation of selection (k<1) in groups marked by K and I and J is tested relative to reference groups marked by G and L respectively (see *Relax: tests for shifts in selection intensity* in Results and Table 3). The **parallel origins/defense escalation model** (subpanels b, d) predicts that the ancestral *HSS*-like locus (Branch A) was under neutral selection (ω=1) followed by multiple independent origins of PAs, and independent optimization of HSS function under positive selection (ω>1) in the ancestral *HSS* loci of PA-producing clades: Echiteae (Branch C) and within Apocyneae (Branches D and E, subpanels b and d), and/or in *HSS* loci in which the PA-associated VXXXD motif evolved. Evolution of the VXXXD motif is reconstructed either in the ancestral HSS locus of Nerieae and Malouetieae (Branch G, subpanel b) or independently in the ancestral HSS loci of *Isonema*, *Strophanthus*, and Malouetieae (Branch F) (subpanel d). VXXXD is also reconstructed as evolving in the ancestral HSS of Apocyneae and MOE (Branch B) in both trees (subpanels b and d). Asterisks at nodes represent bootstrap support > 90%.

**Fig. 4:**
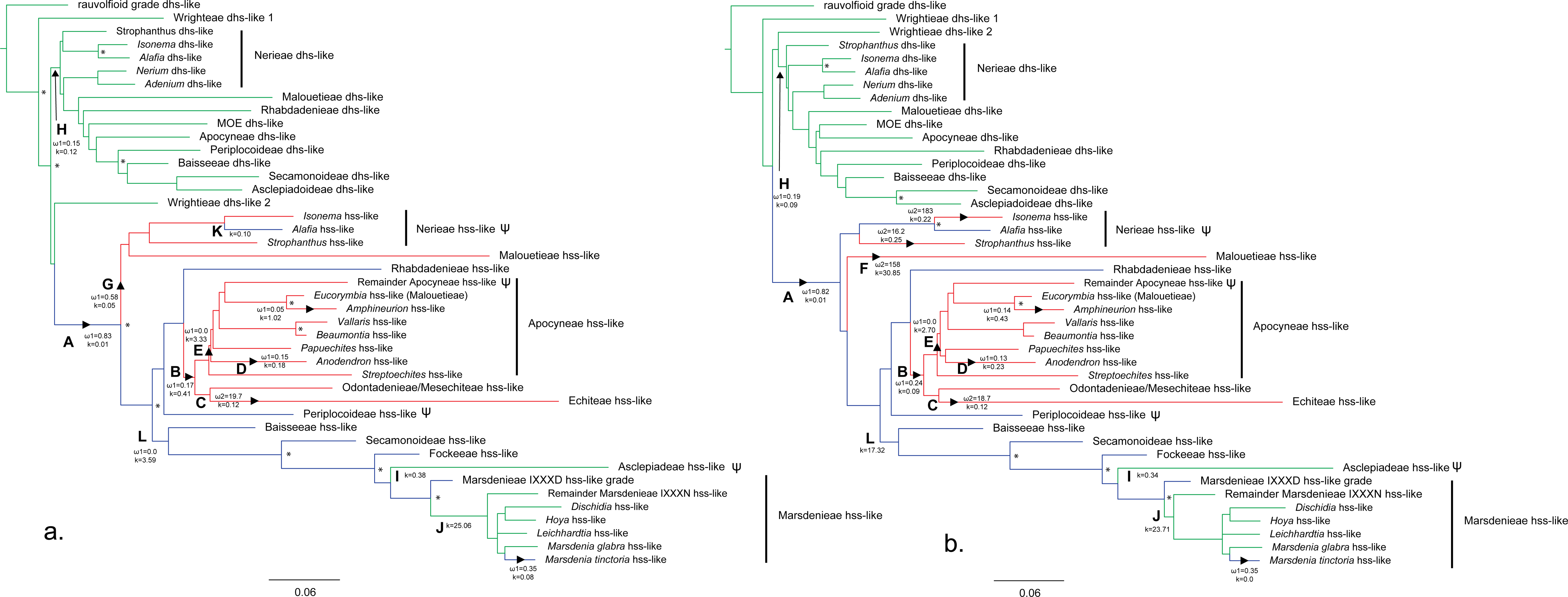
*DHS/HSS* gene trees, collapsed to display lineages of interest (see Methods: *Tree Construction*). **a.** reduced dataset gene tree (Figs. 3a, b for summary gene tree, Appendix S4 for full gene tree); **b**. full dataset topology gene tree (Figs. 3c, d for summary gene tree, Appendix S5 for full gene tree). Branch lengths are proportional to probability of nucleotide substitution. Branch color reflects the reconstructed amino acid motifs (VXXXD in red, IXXXD in blue, IXXXN in green). Branches and lineages A-L, used in selection tests, (see Fig. 3, Tables 1-3) are labeled with letters and arrowheads. aBSREL (ω; Table 1) and RELAX (k; Table 3) results are displayed at selected branches. Bootstrap support > 90 % is indicated with asterisks.

**Table 1.**
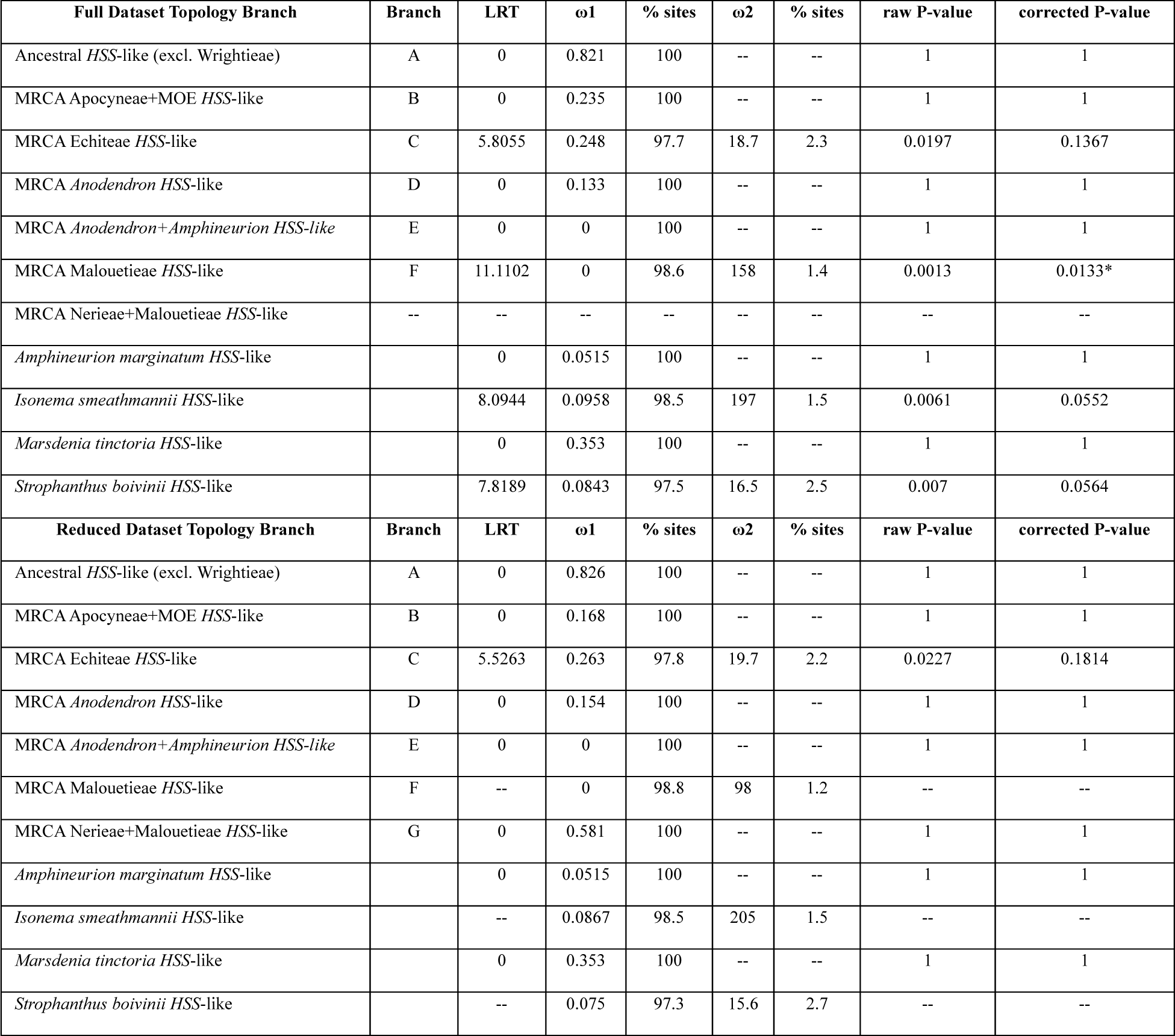
Tests for positive selection on a subset of sites on selected branches in the full dataset and reduced dataset topologies with aBSREL. Branches lettered in Figs. 3 (both topologies), 4 (both topologies), Appendix S4 (Reduced dataset topology) and Appendix S5 (Full dataset topology); “--” indicates that the branch does not occur or was not tested in the topology. The likelihood ratio test (LRT) compares the likelihood of the null (ω <1) versus the alternative model (optimized ω). The optimized model fits one or two ω classes to these data, with >97% of sites evolving under ω1 and <3% of sites evolving under ω2. LRT=0 indicates no improvement in model fit and LRT >>0 indicates better model fit, significance indicated by p-values (raw and corrected for multiple testing). Significant p-values indicated with asterisk.

**Table 2.**
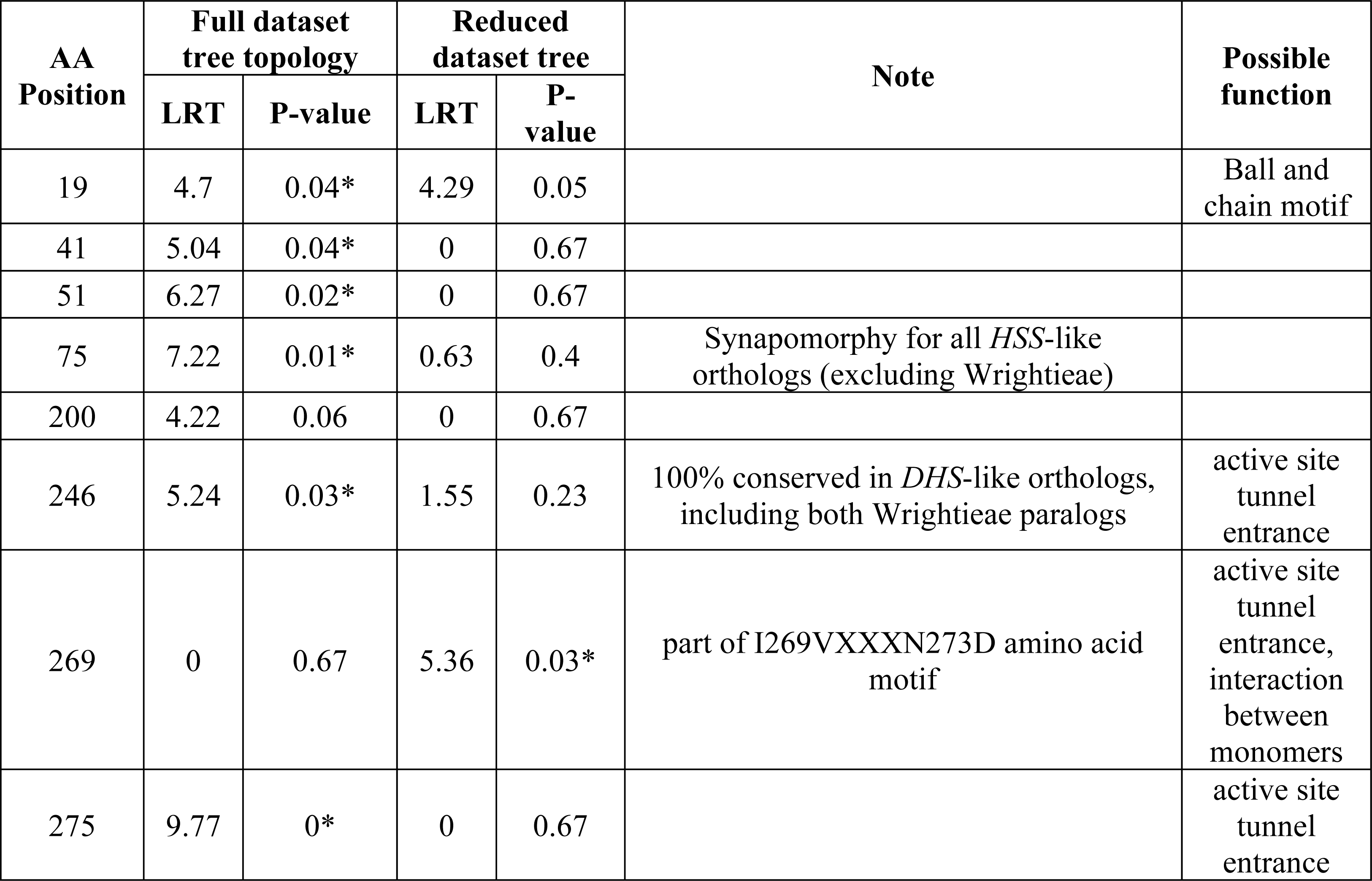
Tests for positive selection on individual amino acid positions on designated branches with MEME. Lettered branches A-G, as well as the *HSS-*like branches of *Amphineurion marginatum*, *Isonema smeathmannii*, *Marsdenia tinctoria*, and *Strophanthus boivinii* (Figs. 3, 4) are tested in the reduced dataset topology (Fig. 4a, Appendix S4) and the full dataset topology (Fig. 4b, Appendix S5). The LRT compares the null model, where nonsynonymous to synonymous mutation ratios are restricted to ω <1, to the alternative model, where ω is unrestricted. Only amino acid (AA) positions that experienced significant positive selection in at least one of the topologies are reported; substitutions at these sites are mapped on the tested topologies (Appendices S4, S5). Significant p-values indicated with an asterisk. Possible function is inferred from homologous human DHS amino acid positions.

**Table 3.**
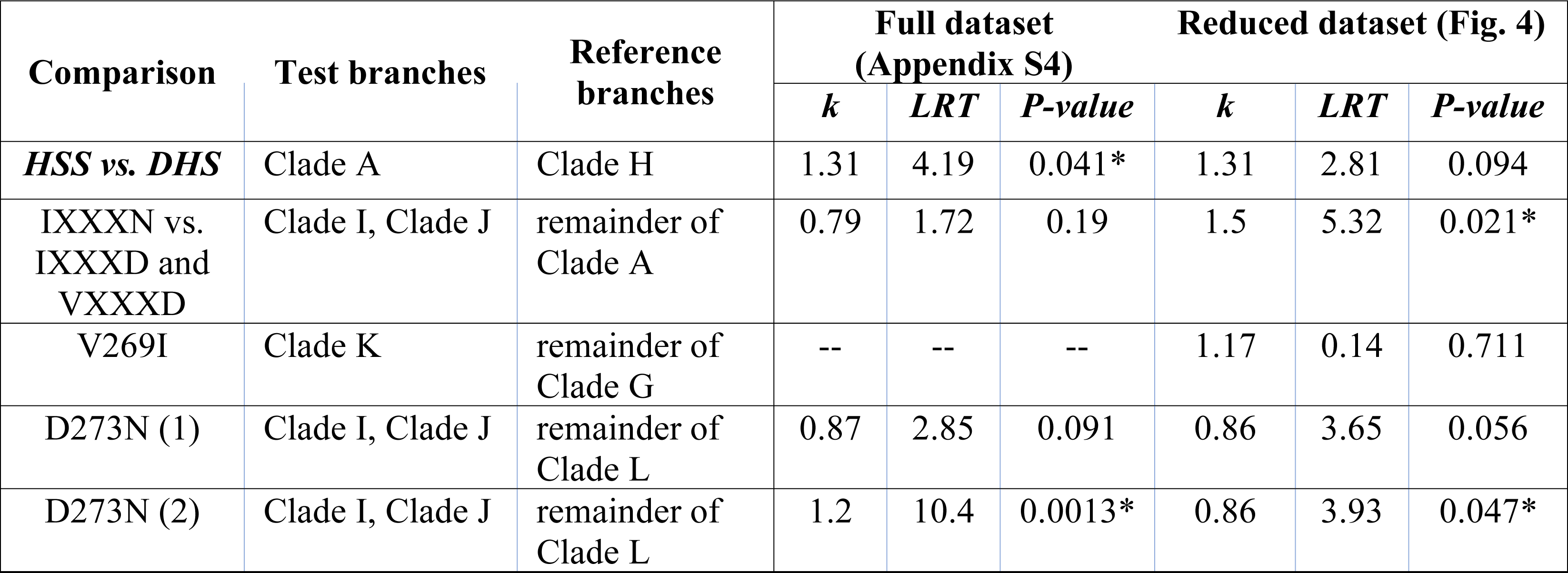
Tests for significant shifts in selection strength on designated branches with RELAX. Branches lettered as in Figs. 3 (both topologies), 4 (both topologies), Appendix S4 (Reduced dataset topology), and Appendix S5 (Full dataset topology). The test branches (i.e. clades subtended by and including the lettered branches) are compared against the reference branches. RELAX estimates k as a selection intensity parameter. The null model fits three ω rate categories to the test and reference branches, and the alternative model raises the rates on the test branches to the power of k, ω^k^. The LRT compares the null and alternative models. A significant k>1 indicates that selection has intensified in test branches relative to reference branches, and a significant k<1 indicates relaxation. Significant p-values (<0.05) indicated by an asterisk. Replicate runs of the D273N test [D273N (1) and D273N (2)] produced inconsistent results.

### Tests for recombination—

Marsdenieae *HSS*-like paralogs were investigated further using GARD (Kosakovsky Pond et al., 2006). GARD searches an alignment for a maximum number of breakpoints, builds phylogenies for every non-recombinant contig, and assesses those phylogenies using Akaike Information Criterion (AIC). Potentially recombinant contigs were split at potential recombination breakpoints indicated by GARD and a Marsdenieae *HSS*-like gene tree (outgroup: *Tassadia propinqua HSS*-like gene) was rebuilt using maximum likelihood tree construction criteria described above.

### Shimodaira-Hasegawa tests—

To test alternate topologies of Apocynaceae *DHS-/HSS*-like gene trees and test the monophyly of IXXXN and IXXXD paralogs in Marsdenieae, Shimodaira-Hasegawa (SH) tests in RAxML-HPC2 on XSEDE (v8.2.12) (Stamatakis, 2014) were performed. The SH test rejects or fails to reject a null hypothesis of equal support for two given topologies.

### Ancestral sequence reconstruction—

Ancestral sequences were reconstructed using codeml Model M0 (default except: model=0, NSsites=0, RateAncestor=1, cleandata=0) in PAML v4.9j (Yang, 1997; Yang, 2007). codeML integrates the Goldman and Yang model of amino acid substitution and assumes that selection pressure on an individual site is the same for every branch, produces joint likelihood reconstructions (all ancestral nodes reconstructed), and uses empirical Bayes procedure (Yang and Wang, 1995) for sequence reconstruction. Additionally, codeml calculates a marginal reconstruction (single nodes reconstructed), which includes posterior probabilities for reconstructed amino acids.

### Tests for selection: hypotheses—

If origin of pyrrolizidine alkaloids was adaptive, and if selection for PAs caused adaptive evolution of *HSS*, we can make testable predictions about patterns of selection on *HSS*-like genes in Apocynaceae. If there was a single origin of PAs in the MRCA of all PA-producing taxa (Figs. 3a, c), we predict positive selection on the *HSS*-like gene of this MRCA (Figs 3a, c, branch A, ω>1) followed by purifying selection on branches that retained optimized HSS function (i.e. IXXXD motif in clade L or VXXXD motif in clade G) and relaxed selection on branches that lost optimized HSS function (i.e. evolution of IXXXN motif, Figs. 3a, c, clades I, J, ω=1, k<1). We also test if the evolution of an IXXXD (Fig. 3a, clade K, ω=1, k<1) in two PA-free *Alafia* species from an ancestral VXXXD motif (Fig. 3a, clade G) is a result of loss of function. [While PAs have been reported from *Alafia*, the two sequenced species tested negative for PAs (Barny et al., 2021).] In contrast, if parallel recruitment of the ancestral *HSS*-like paralog into independently evolved PA biosynthetic pathways occurred (Figs. 3b, d), we would predict relaxed selection (due to loss of DHS function) on the ancestral *HSS*-like gene (Figs 3b, d, branch A, ω=1) followed by positive selection on the ancestral *HSS* of each lineage were the HSS VXXXD motif (present in all sequenced PA-producing Apocynaceae species) and/or PA-production evolved (Figs 3b, d, branches B, C, D, E, F, G, to *Isonema*, to *Strophanthus*, ω>1). Under this scenario, branches with IXXXN and IXXXD motifs should remain under relaxed selection (if they did not evolve some new function); we predict relaxation of selection in the HSS clade (clade A, k <1) relative to the DHS clade (clade H), since most HSS branches would be non-functional and under relaxed selection (Figs 3b, d).

### Tests for selection: analyses—

[A]BSREL (adaptive Branch-Site Random Effects Likelihood) (Smith et al., 2015) was used to test for positive selection on selected branches (Fig. 3, ω>1). It fits optimal ω (dN/dS ratio) distributions to each branch by assigning each site to one of up to three ω rate categories. This generates the optimal ω distribution for each branch. For branches with more than one ω rate, the larger one is mapped in Fig. 4, Appendices S4 and S5. Positive selection is inferred on *a priori* selected branches by comparing, via likelihood ratio test (LRT), the optimized ω distribution to a null model with constraint ω<1 for all sites.

MEME (Mixed Effects Model of Evolution) (Murrell et al., 2012) was used to identify sites under positive selection on pre-specified branches (Fig. 3, ω>1). It allows selection to vary both among branches and among sites. Each branch is assigned one of two ω rate classes at each amino acid site. A single α (synonymous substitution rate) is shared among all branches. First nonsynonymous substitution rate (β-) is estimated for each site; β-is constrained to be less than or equal to α (i.e. evolving neutrally). Second nonsynonymous substitution rate (β+) is unconstrained in the full model. Likelihood ratio tests are used to compare the full model with a null model where β+ is constrained to be less than α.

RELAX (Wertheim et al., 2015) was used to test for neutral evolution. It tests for relaxation and intensification of positive and purifying selection on pre-specified test branches (Figs 3a, c, lineages K, I, J) compared to a set of designated reference branches (Figs 3a, c, lineages G, L). The RELAX null model assigns all sites into one of three rate classes (ω1=purifying, ω2=neutral, ω3=positive selection). The full (alternative) model introduces a selection intensity parameter, k, and raises ω^k^ on the test branches. The null model constrains k=1, which forces the same ω distribution in both the test and reference branch sets. If the likelihood ratio tests find the alternative model is significantly better fit, a k>1 is considered evidence of intensified selection and a k<1 relaxed selection in the test branches relative to the designated reference branches. The RELAX general descriptive model was used to calculate the k values mapped in Fig. 4, Appendices S4 and S5, from RELAX analyses comparing all *HSS*-like branches to *DHS*-like branches (Appendix S1, Table S12.4). Rather than using the *a priori* test and reference branch sets, the general descriptive model fits the three ω rates to all branches, and an individual k for each branch (Wertheim et al., 2015).

### Comparison with human DHS—

The reduced dataset was aligned with the angiosperm *DHS/HSS* alignment from Livshultz, et al. (2018) and human *DHS* (Genbank: P49366) to produce a “human and plant *DHS/HSS* alignment” (Appendix S1, Table S3). Human DHS amino acid site functions are described in Wator, et al. (2020). The amino acid positions of DHS monomer interaction sites and general functional sites (i.e. active site tunnel entrance) were inferred from annotations on structures PDB ID 6XXXM using the NCBI Structure feature (Madej, et al. 2014) on iCN3D v.2.24.4 (Wang, et al. 2020). These sites were manually annotated on the human *DHS* sequences in the alignment to enable comparisons between human and plant DHS/HSS amino acid positions.

### Site-directed mutagenesis Parsonsia alboflavescens HSS—

The open reading frame of *Parsonsia alboflavescens* HSS (*Pa*HSS), cloned in an expression vector (NovagenTM pET28a, Millipore Sigma, Billerica, MA, USA) with an artificial N-terminal hexahistidine (6xHis) tag extension, was used as template for site-directed mutagenesis guided by Liu and Naismith (2008). Primer pairs to introduce the single mutations V269 to I269 (numbering of the amino acids follows Kaltenegger, et al. 2013) and D273 to N273 as well as to double mutation V269XXXD273 to I269XXXN273 are given in Appendix S1, Table S4. PCR amplifications were performed in 25 µl reaction volume with Phusion® High-Fidelity DNA Polymerase (Thermo-Scientific) according to the manufacturer’s instructions; annealing temperature is given in Appendix S1, Table S4. Twelve (12) amplification cycles were performed. The PCR products were treated with *Dpn*I (Thermo-Scientific) at 37°C for 1 hour, diluted with water (1:10), subsequently propagated in *Escherichia coli* TOP10 (Thermo-Scientific), and sent out for Sanger sequencing (MWG Eurofins Genomics) to identify successful mutants.

### Heterologous expression, purification and activity assays of *P. alboflavescens* HSS and mutants—

The complete ORF of the *Pa*HSS and the mutant variants were expressed in *Escherichia coli* BL21(DE3) and purified according to (Ober and Hartmann, 1999). Protein purification was monitored via SDS-PAGE analysis and protein quantities were estimated based on UV absorption at 280 nm and the specific extinction of the respective protein, calculated with the PROTPARAM web tool on ExPASy (Gasteiger et al., 2005) and with the Bradford method (Bradford, 1976). The oligomerization state of the purified proteins was analysed by size exclusion chromatography coupled to UV. Eight (8)-15 µg of affinity-purified DHS and HSS in borate buffer (∼ 42 kDa) were analyzed on an analytical size-exclusion chromatography (SEC) column (MabPac Sec-1 5 µm 300 Å, 4 x 150 mm) equilibrated with 50 mM phosphate buffer (pH 6.8) plus 0.3 M NaCl (0.2 ml/min flow), connected to an Ulitmate 3000 system and a DAD-3000 diode array detector (Thermo Fischer Scientific, Waltham, MA, USA). Proteins were monitored at 280 nm. Cytochrome C (12 kDa) and BSA (monomer 66.5 kDa, dimer 132 kDa) were used as reference proteins.

For biochemical characterization, the purified proteins were concentrated and rebuffered to borate-based (50 mM borate-NaOH buffer, pH 9) assay buffer which included the additives DTT (1 mM) and EDTA (0.1 mM). The *in vitro* assays were performed according to (Kaltenegger et al., 2021). In short, 5 – 40 µg purified recombinant protein were incubated with putrescine and spermidine (400 µM each), in the presence of NAD (2 mM), in borate -based assay buffer to determine the enzyme’s ability to produce homospermidine. Product formation was quantified via derivatizing the reaction mixture with 9-fluorenylmethyl chloroformate (FMOC, Sigma) and subsequent analyses by HPLC coupled with UV detection. To detect the enzymes’ ability to utilize the eIF5A, assays were hydrolyzed as described in Kaltenegger, et al. (2021) derivatized with FMOC and analysed by HPLC coupled with FLD to quantify deoxyhypusine, 1,3-diaminopropane and canavalmine.

## RESULTS

### Taxon sampling—

We obtained *DHS*-like and *HSS*-like sequences from 142 species in 14 of 16 APSA clade tribes or subfamilies (not sampled: Ceropegieae, Eustegieae) and 9 of 11 rauvolfioid tribes (not sampled: Alstonieae, Willughbeieae) (Fig. 1), from new sequencing and published datasets (Hoopes et al., 2018; Livshultz et al., 2018) (Appendix S1, Table S2). Taxonomy follows Endress et al. (2018).

### Assembly and annotation of DHS- and HSS-like loci—

Our automated assembly pipeline yielded 437 annotatable contigs (Appendix S1, Table S2) from 122 libraries, 113 of which contained a complete CDS. Contigs with a partial ORF were manually concatenated according to a topology-aware algorithm (see ‘gene assembly from contigs’ in Materials & Methods). After concatenation, we recovered at least one *DHS*-like gene with an intact ORF covering at least exons 2 through 6 from all 122 libraries (Appendix S1, Table S2); all rauvolfioid species, Wrightieae species, and the outgroup (*Gelsemium sempervirens*) yielded only *DHS*-like contigs. There was at least one *HSS*-like gene, defined as an ortholog of *Parsonsia alboflavescens HSS*, with an intact ORF covering at least exons 2 through 6 from 87 libraries. All APSA clade species, except Wrightieae, had *HSS*-like contigs, either with intact ORFs or internal stop codons.

Twenty-nine (29) accessions previously sequenced by Livshultz et al. (2018) with the Sanger method were re-sequenced and assembled (Appendix S1, Table S2). All 36 re-sequenced loci have over 96% amino acid identity in the region of overlap with the Sanger sequences. We also found 43 additional sequences from these 29 libraries: full *DHS*-like sequences in 11 libraries, full *HSS*-like sequences in 6 libraries, short *DHS*- or *HSS*-like sequences in six libraries, and *DHS*- or *HSS*-like pseudogene sequences in 8 libraries.

### Distribution of DHS and HSS motifs—

All *DHS*-like sequences with intact ORFs encode the expected IXXXN motif at positions 269 and 273 except *Strophanthus boivinii* (MXXXN) (Appendix S1, Table S2). *HSS*-like sequences with intact ORFs encode IXXXN, IXXXD, and VXXXD motifs. Amino acid motifs in translated *HSS*-like sequences are conserved at the tribal/subfamilial level. *HSS*-like genes in Malouetieae, Apocyneae, Mesechiteae, Odontadenieae, and Echiteae encode a VXXXD motif (Fig. 1). Sequences from Secamonoideae, Periplocoideae, Baisseeae, Fockeeae, and Rhabdadenieae encode IXXXD, and Asclepiadeae sequences, IXXXN (Fig. 1). Nerieae and Marsdenieae have multiple motifs. Among Nerieae, *Strophanthus* and *Isonema* have VXXXD, while *Alafia* has IXXXD (Figs. 1, 3, 4). Marsdenieae *HSS*-like sequences encode IXXXD or IXXXN, with one exception, *Dischidia albida* (MXXXT) (Fig. 1, Appendix S6). All 10 sequenced PA-producing Apocynaceae species have an intact VXXXD *HSS*-like gene (Appendix S1, Table S2).

### Pseudogenes—

Fifty-six (56) putative pseudogenes, i.e. sequence assemblies with internal stop codons, occurred in both the *DHS*-like grade and the *HSS*-like clade (Appendix S1, Tables S2, S3; Appendix S3). No species had both *DHS*-like and *HSS*-like pseudogenes. In most cases, these putative pseudogenes co-occurred with intact *DHS*- or *HSS*-like genes. However, nine species in five APSA lineages have only *HSS*-like pseudogenes: Apocyneae (*Apocynum androsaemifolium, A. cannabinum, A. pictum, A. venetum* [two accessions]), Asclepiadeae (*Asclepias syriaca),* Malouetieae (*Pachypodium baronii*), Nerieae (*Nerium oleander, Strophanthus preussii*), and Periplocoideae (*Zygostelma benthamii*) (Figs. 1, 4, Appendix S3).

### Final alignments and gene tree topologies—

The final alignments contained sequences with an intact ORF covering at least exons 2-6, which includes 120 *HSS*-like and 155 *DHS*-like sequences, 238 of them new (Appendix S1, Table S2). The “full dataset” contained additional shorter sequences and pseudogenes, which were both excluded in the “reduced dataset” (Appendix S1, Table S3).

Topologies of the maximum likelihood trees from the full dataset (Figs. 3c, d, 4b, Appendix S5) and the reduced dataset (Figs. 3a, b, 4a, Appendix S4) were very similar. The most important differences are the position of the Wrightieae *DHS*-like 2 clade, sister to the *HSS*-like clade (Branch A) in the reduced dataset tree (Figs. 3a, b, 4a, Appendix S4) or sister to the APSA *DHS*-like clade in the full dataset tree (Figs. 3c, d, 4b, Appendix S5), and the monophyly of Nerieae plus Malouetieae *HSS*-like genes in the reduced dataset tree (Branch G, Figs. 3a, b, 4a, Appendix S4) or their paraphyly in the full dataset (Figs. 3c, d, 4b, Appendix S5). A Shimodaira-Hasegawa (SH) test could not reject these alternate topologies (Appendix S1, Table S6) so both topologies were used in downstream analyses. Because bootstrap support values were higher in the reduced dataset tree (Fig. 4a, Appendix S4) relative to the full dataset tree (Fig. 4b, Appendix S5), gene tree topology will be discussed with reference to the reduced dataset tree (Figs. 3c, d, 4a, Appendix S4) except when noted.

All APSA clade sequences form a highly supported monophyletic clade (98 BS), nested in a grade of rauvolfioid sequences (Fig. 4a, Appendix S4). Within the APSA clade, HSS-like sequences are in a fully supported clade (Branch A, Fig. 4a, Appendix S4). Clades of sequences from related taxa (classified in the same genus, tribe, or subfamily) are monophyletic and mostly moderately to well-supported (Fig. 4a, Appendix S4). Exceptions include tribes and genera whose monophyly has not been consistently supported in previous phylogenetic analyses (Odontadenieae, Nerieae, *Marsdenia*) (Livshultz et al., 2007; Morales et al., 2017; Fishbein et al., 2018; Espírito Santo et al., 2019; Liede-Schumann et al., 2022), as well as the position of *Eucorymbia alba* (Malouetieae) as sister to *Amphineurion marginatum* (100 BS *HSS*-like clade, 98 BS *DHS*-like clade) in the Apocyneae, a placement likely reflecting its phylogeny. Relationships among tribal and subfamilial gene lineages also reflect species phylogeny (Fig. 1). In both *HSS*-like and *DHS*-like subtrees, Wrightieae, Nerieae, and Malouetieae sequences diverge prior to the “crown clade” radiation comprising the rest of the APSA clade (Straub et al., 2014; Fishbein et al., 2018; Antonelli et al., 2021); Baisseeae are sister to the Secamonoideae and Asclepiadoideae clade (Fockeeae, Asclepiadeae, Marsdenieae) (Antonelli et al., 2021). Sequences from the predominantly New World Mesechiteae, Odontadenieae, and Echiteae form the “MOE” clade (83 BS *HSS*-like clade, 88 BS *DHS*-like clade) (Livshultz et al., 2007; Morales et al., 2017). The incongruent positions of “MOE”, Apocyneae, Rhabdadenieae, and Periplocoideae between the *DHS*- and *HSS*-like subtrees likely reflect long branch attraction and/or incomplete lineage sorting in a known rapid radiation (Straub et al., 2014).

Within the APSA HSS-like clade, at least one duplication gave rise to the paralogous IXXXN and IXXXD *HSS*-like loci found in 13 of 23 Marsdenieae species. While the maximum likelihood topology resolved the IXXXD *HSS*-like loci as a grade forming successive sister lineages to an unsupported IXXXN *HSS*-like clade (Clade J; Figs. 3, 4; Appendices S4, S5), an SH test could not reject the alterative topology with the grade-forming IXXXD *HSS*-like sequences instead forming a monophyletic clade (Appendix S1, Table S6). Motifs were homogenous in these two gene lineages except for a *Marsdenia tinctoria* locus (IXXXD) and *Dischidia albida* (MXXXT) locus nested in the IXXXN *HSS*-like clade and the *Gongronemopsis truncata HSS2* sequence (IXXXN) in the IXXXD *HSS*-like grade (Appendix S6, a). These sequences were assembled by the pipeline without manual concatenation and read mapping did not reveal assembly errors. When sequences were split at a recombination break point indicated by GARD (Kosakovsky Pond et al., 2006) and realigned, it was determined that, although the *Gongronemopsis truncata* (IXXXN) sequence is recombinant, the discordant placement of its amino acid motif is not a product of recombination (Appendix S6, b). The *Marsdenia tinctoria* sequence (IXXXD) is not a recombinant since both the 5’ and 3’ ends of the sequence nest within the IXXXN clade (Appendix S6, b).

### Ancestral motif reconstruction—

To reconstruct ancestral motif at amino acid positions 269 and 273, we reconstructed evolution of the reduced dataset alignment on both the reduced dataset tree topology (Figs. 3a, b, 4a, Appendix S4) and a topology constrained to match the full dataset topology (Figs. 3c, d, 4b, Appendix S5) with both joint and marginal (Appendix S1: Tables S7, S8) likelihood in PAML (Yang, 1997; Yang, 2007). The two methods gave highest likelihood to the same reconstructed motifs (Appendix S1: Tables S7, S8), hence the marginal reconstruction will be discussed.

On the full dataset tree topology (Figs. 3c, d, 4b, Appendix S5), the ancestral motif for the *HSS*-like subtree was IXXXD (Branch A). VXXXD motifs evolved four times from IXXXD motifs: 1) *Strophanthus boivinii*, 2) *Isonema smeathmannii*, 3) MRCA Malouetieae (Branch F), and 4) MRCA of Apocyneae+MOE (Branch B). There were also three substitutions from IXXXD to IXXXN, a reverse mutation back to the DHS-like motif: 1) in the MRCA of the Asclepiadeae tribe (Branch I), 2) in the MRCA of the Marsdenieae IXXXN *HSS*-like clade (Branch J), and 3) in *Gongronemopsis truncata HSS-*like 2. A single re-evolution of IXXXD from IXXXN occurred in the *Marsdenia tinctoria HSS-*like gene.

The reduced dataset tree topology (Figs. 3a, b, 4a, Appendix S4) differed only in that a VXXXD motif evolved only twice from the ancestral HSS-like IXXXD motif: in the MRCA of Apocyneae+MOE (Branch B) and in the MRCA of Nerieae-Malouetieae (Branch G). From a VXXXD there was one reversal to IXXXD in the MRCA of the two *Alafia* species (Branch K).

### Evidence for selection on HSS loci—

We used aBSREL (Smith et al., 2015), which tests if a proportion of sites in specified branches evolved under positive selection, and MEME (Murrell et al., 2012), which tests for site-specific positive selection, to test our main hypotheses: single (Figs. 3a, c) versus multiple convergent origins (Fig. 3b, d) of optimized HSS and PA biosynthesis. Selection criteria for tested branches are 1) evolution toward an HSS-like motif (I269V and/or N273D), or 2) occurrence of PAs (Fig 3, branches A, B, C, D, E, F, G).

Significant positive selection, after correction for multiple testing, was only found on the MRCA of Malouetieae (Branch F) in the full dataset topology tree (Figs. 3c, d, 4b, Appendix S4, Table 1). There was no evidence of significant positive selection on the ancestral HSS-like branch (Branch A) (Fig. 3, 4, Appendices S4 and S5, Table 1), nor for any branches where a more HSS-like motif evolved or where PAs occur (Table 1).

We hypothesized that individual sites experienced positive selection on branches where optimized HSS function evolved. For the full dataset tree topology (Figs. 3c, d, 4b, Appendix S5), six amino acid sites showed evidence of significant positive selection on these branches: sites 19, 41, 51, 75, 246, and 275 (Table 2). For the reduced dataset tree (Figs. 3a, b, 4a, Appendix S4), only one amino acid site showed evidence of significant diversifying selection on the test branches: 269 (Table 2).

Position 269, importantly, is the first amino acid in the focal motif (I/VXXXN/D). Of the remaining seven sites, two may be important for optimization of HSS function and/or loss of eIF5A activation based on substitution patterns. Position 75 is a synapomorphy for the *HSS*-like clade (D75A substitution on Branch A) in both trees (Appendices S4 and S5). Position 246, which is adjacent to an active site position in human DHS (Wator et al., 2020), is 100% conserved in *DHS*-like sequences, including both Wrightieae paralogs. However, L246I substitutions occur 7 times in the *HSS*-like clade: on the branches leading to *Isonema smeathmannii* (Nerieae), *Galactophora schomburgkiana* (Malouetieae), the MRCA of Echiteae (Branch C), the MRCA of *Amphineurion*+*Eucorymbia* (Apocyneae), the MRCA of Asclepiadeae (Branch I), the MRCA of Periplocoideae, and the MRCA of the *Mandevilla HSS*-like clade (Figs. 3, 4; Appendices S4 and S5). This same substitution, L246I, is present in all other functionally characterized angiosperm HSS except *Eupatorium cannabinum, Ipomoea alba, I. hederifolia, and I. neei* (Appendix S1, Table S3).

### Tests for shifts in selection intensity—

We used RELAX, which detects changes of selection strength (Wertheim et al., 2015), to test our hypothesis that evolution of a more DHS-like motif from a more HSS-like motif in *HSS*-like genes is evidence of loss of PA biosynthesis, accompanied by relaxation of selection on HSS function and a shift toward neutral evolution (as predicted by the single origin, defense de-escalation model, Figs. 3a, c). Test clades where a more DHS-like motif (IXXXN or IXXXD) evolved from a more HSS-like motif (IXXXD or VXXXD) (Figs 3a, c, clades I, J, K) were compared to reference clades which retained the more HSS-like motif (Figs 3a, c, clades G, L). None of the predictions derived from the defense de-escalation hypothesis were supported (Table 3). In testing IXXXD branches (Clade K) against VXXXD branches in Clade G, which only occurred in the reduced dataset topology (Figs. 3a, 4a, Appendix S4), the RELAX null model was a better fit than the alternative model, supporting no shift in selection intensity (Appendix S1, Table S12.1). In testing IXXXN branches (Clade I, Clade J, *Gongronemopsis truncata HSS*-like 2) against reference IXXXD branches in Clade L (Figs. 3a, c), the results differed between tree topologies, and replicate runs for both tree topologies had variable results, issues with convergence (Table 3; Appendix S1: Table S12.2) and warnings that results were untrustworthy. Similarly, in testing all IXXXN *HSS*-like branches (Figs 3a, c, clades I, J) against all IXXXD and VXXXD *HSS*-like branches (Clade A, Figs. 3a, c), results were not consistent between tree topologies (Table 3; Appendix S1: Table S12.3). This variation makes interpretation of selection intensity on IXXXN vs IXXXD/VXXXD branches impossible.

Lastly, we hypothesized that the APSA *HSS*-like genes (clade A) are under relaxed selection relative to *DHS*-like genes (clade H) because DHS function is essential in primary metabolism while the rare occurrence of PAs predicts that most *HSS*-like genes are without function (Fig. 3b, d). The alternative model for both tree topologies was significantly better than the null model, supporting an intensification of positive selection on the *HSS*-like clade (Appendix 1, Table S12.4). This intensification of selection was significant in the full dataset topology but not in the reduced dataset (LR = 2.81, P = 0.094). This can be interpreted as an overall stronger signal for positive selection for optimized HSS function than relaxation of selection on HSS function when averaged across all branches of the *HSS* subtree.

There is a pattern of relaxation of selection in ancestral *DHS*-like and *HSS*-like branches, as illustrated by the optimal k-parameters inferred by the RELAX general descriptive model (Figs. S3, S4). After the core APSA *DHS/HSS* duplication event, there was a relaxation of purifying selection on the ancestral *DHS*-like Branch H (k=0.09 to 0.12) and the ancestral *HSS*-like Branch A (k=0.01) (Fig. 4, Appendices S4, S5; Appendix 1: Tables S10, S11). The succeeding three branches in the backbone of the core *DHS*-like clade then experienced intensification of selection, while most of the backbone of the core *HSS*-like clade continued to experience predominantly relaxation of selection (Appendices S4, S5; Appendix 1: Tables S10, S11).

### Function of HSS VXXXD motif—

To test the functional relevance of the VXXXD motif in Apocynaceae HSS, we constructed single (IXXXD, VXXXN) and double (IXXXN) mutants of the HSS from *Parsonsia alboflavescens* [a PA producing species (Barny et al., 2021)] and tested their activity with putrescine to produce homospermidine (HSS function, Fig. 2b) and the eIF5A precursor protein to produce activated eIF5A with a lysine aminobutylated to deoxyhypusine (DHS function, Fig. 2a) in *in vitro* assays. All four enzymes were successfully expressed and purified (Appendix S7). SEC-UV confirmed that the expressed enzymes assembled to a tetramer, the biologically active unit of HSS (Umland et al., 2004; Wator et al., 2020).

To compare HSS activity among the enzymes, the specific activity for homospermidine production was calculated (Table 4). The activity of the IXXXD single mutant is comparable to that of wild type VXXXD HSS (mean 1145±34.35 pkat/mg, 1317±210.72 pkat/mg, respectively). Both the VXXXN single (mean 471±4.71 pkat/mg) and the IXXXN double (mean 526±36.82 pkat/mg) mutants show a decreased specific activity compared to the wild-type enzyme. 1,3-diaminopropane (Dap) was produced in roughly equimolar amounts with homospermidine, indicating that the transfer of the aminobutyl moiety to putrescine was the major reaction catalyzed by the enzymes. HSS activity is high in all four enzymes when compared to *P. alboflavescens* DHS, 274 pkat/mg (Livshultz et al., 2018). The two-fold higher activity measured for the IXXXD and VXXXD enzymes compared to the VXXXN and IXXXN enzymes may be due to loss of catalytic activity caused by the D273N substitution (suggesting functional importance of this substitution for HSS optimization) or to a greater ratio of inactive to active enzyme in the latter two enzyme preparations (suggesting an experimental artifact).

**Table 4.**
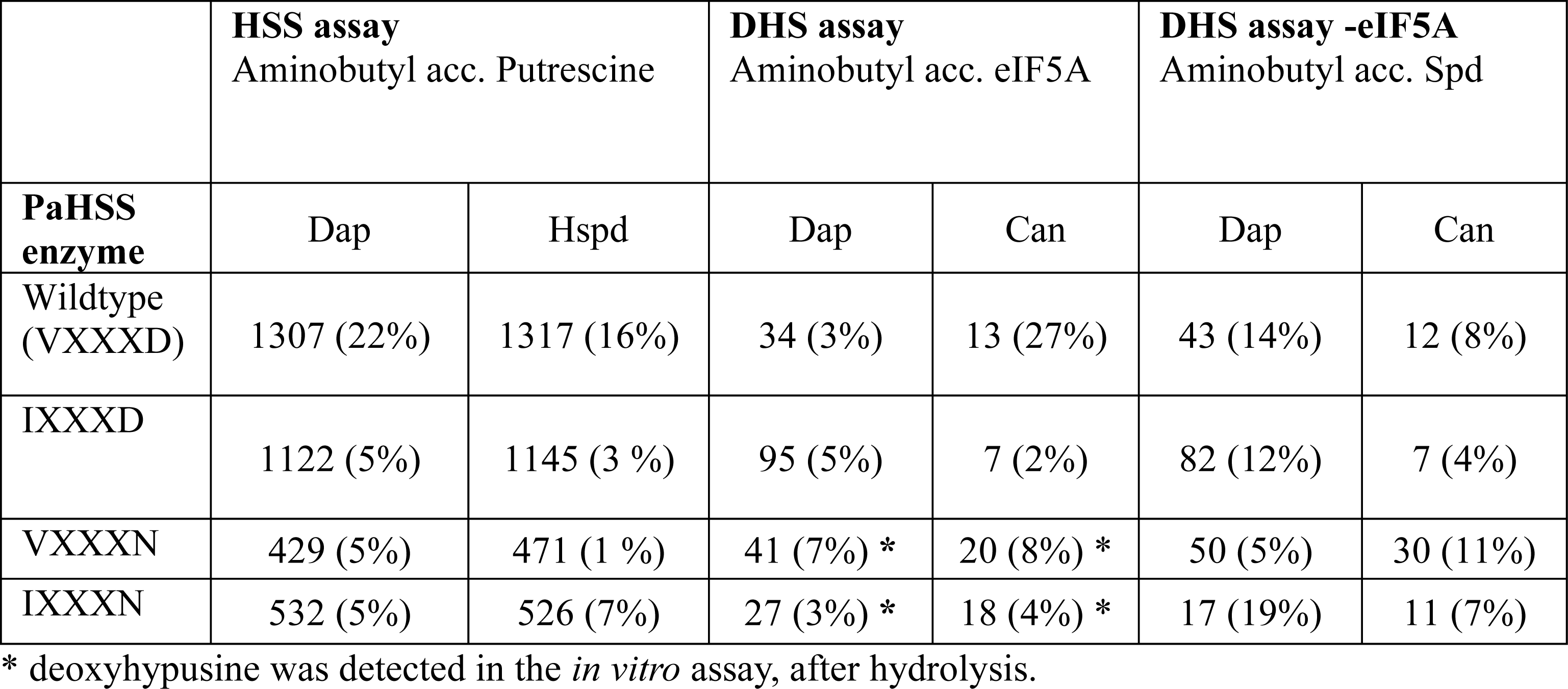
Mean specific activities of wild-type and mutated *Parsonsia alboflavescens* HSS. For the HSS function assay (substrate=putrescine), the specific activity of homospermidine (Hspd) production, as well as 1,3-diaminopropane (Dap) was measured. Both DHS function assays (substrate=eIF5A and substrate=spermidine [Spd]) measured production of Dap and canavalmine (Can), Values in pkat mg^-1^ and the relative standard deviation of three replicates are reported in parentheses.

To assess DHS activity, Dap formation was quantified in the presence of eIF5A precursor protein and compared to a negative control assay without it (Table 4). Wildtype (VXXXD) and the IXXXD mutant produced similar amounts of Dap and canavalmine (Can) in both assays, indicating that both enzymes do not aminobutylate the eIF5A precursor protein (Table 4, Appendix S8) but instead use spermidine (Spd) as aminobutyl acceptor (Fig. 2c, reaction II). Of note, while Can increased constantly over time, Dap was produced in a burst at the beginning of the reaction, most evident in the IXXXD mutant (Appendix S8). This burst most likely results from Spd cleavage to Dap and delta pyrrolinium, without transfer of the aminobutyl moiety (Fig. 2c, reaction III). The VXXXN mutant in the assay with the eIF5A precursor produced slightly less Dap (mean 41±3.29 pkat/mg) than when eIF5A precursor was excluded (mean 50±2.5 pkat/mg) (Table 4, Appendix S8). In contrast, the double mutant IXXXN produced more Dap and Can when eIF5A was included (Table 4, Appendix S8). These difference in the DHS assay with versus without eIF5A prompted us to screen for deoxyhypusine in the hydrolyzed assays. Of note, deoxyhypusine, direct evidence of DHS function, could only be detected when VXXXN and IXXXN but not VXXXD or IXXXD enzymes were incubated with the eIF5A precursor protein (Table 4). Thus, the N273D substitution might be important for interaction of the enzyme with eIF5A. It also affects the rate of spermidine cleavage, which is highest in the IXXXD mutant (Appendix S8).

## DISCUSSION

Recruitment of paralogous gene copies to the biosynthetic pathways of identical specialized metabolites shared by distantly related taxa is strong evidence of parallel phenotypic evolution (Pichersky and Lewinsohn, 2011). Biosynthesis of a class of metabolites by orthologous genes is consistent with homology of these metabolites shared by related lineages but does not exclude parallel recruitment to independently evolved pathways. Livshultz, et al. (2018) showed that homospermidine synthase (*HSS*), the first gene of the pyrrolizidine alkaloid (PA) biosynthetic pathway, evolved via a single gene duplication in an early ancestor of the APSA clade of Apocynaceae (Fig. 1). Furthermore, they reconstructed an amino acid motif (VXXXD), diagnostic of a functional HSS in extant PA-producing species, in this ancestor. This pattern is consistent with homology of PAs among the few APSA lineages where they occur (Fig. 1, orange circles) and with a co-evolutionary hypothesis to explain both the scattered distribution of PAs among Apocynaceae and the origins of an unusual behavior in their specialist herbivores (Danainae, milkweed and clearwing butterflies) to acquire PAs as adults rather than larvae (adult pharmacophagy).

In the present study, we revisit evolution of *HSS*-like genes with an expanded dataset and ask if correlation of gene sequence, function and phenotype, patterns of functional evolution, and selection on *HSS*-like genes in Apocynaceae are more consistent with a single origin and multiple losses of PA biosynthesis (Fig 1, black star, black Xs) or multiple parallel origins (Fig 1, white stars). We conclude that the evolutionary history of PAs cannot be confidently inferred from evolution of *HSS*-like genes because of uncertainty in the *HSS* genotype to PA phenotype map (Weiss and Fullerton, 2000; Sholtis and Weiss, 2005). We identify future research that can reduce this uncertainty.

### Partial disconnect amonh HSS motif, catalytic function, and PA phenotype limits application of motif evolution to infer phenotypic homology—

To distinguish between homology and parallel evolution of a class of metabolites synthesized by orthologous genes, we need to establish tight correlations among phenotype, gene function, and gene sequence. Then, when evolution of sequence is reconstructed on a gene tree, we can reconstruct functional and, after mapping the gene tree to a species tree, phenotypic evolution. Theoretical considerations suggest that this could be obtained in the case of HSS and PAs since accumulation of abundant precursor amines (e.g. homospermidine) is considered an essential early step in evolution of a novel alkaloid biosynthetic pathway (Lichman, 2020). This can be accomplished by evolution of higher specific activity and/or expression levels of HSS.

Observational evidence suggests that sequence (VXXXD motif) and phenotype are tightly linked in the case of HSS and PA biosynthesis. We have sequenced HSS from 10 PA-producing Apocynaceae species from two of three PA producing lineages (Echiteae and Apocyneae, Fig. 1, Appendix S1, Table S2); all have the VXXXD motif, as do 15 of the 16 other PA-producing angiosperm species that have been genotyped to date (Kaltenegger et al., 2013; Gill et al., 2018; Livshultz et al., 2018; Prakashrao et al., 2022), with the notable occurrence of infraspecific polymorphisms in two species discussed below. *Heliotropium indicum* (AXXXD) is the one exception (Livshultz et al., 2018). An outstanding question is the HSS motif of PA-producing species of *Alafia,* the only known PA-producing genus of Nerieae (Fig. 1). The two species sequenced here have IXXXD motifs (Appendix S1, Table S2) and no detectable PAs in their leaves (Barny et al., 2021). Finally, the observed association is based on a very small sample since the HSS motifs of an estimated 6,000 PA-producing angiosperm species (Kaltner et al., 2020) are unknown.

In addition to correlated evolution of the HSS VXXXD motif and PAs across angiosperms (Livshultz et al., 2018), natural infra-specific polymorphisms point to the functional importance of the HSS VXXXD motif. In *Lolium perenne* L. (Poaceae), with two HSS loci: *LpHSS1* (VXXXD) and *LpHSS2* (VXXXN), plants homozygous for the null allele at the *LpHSS1* locus produce ca. 5 times less PAs than plants with at least one functional *LpHSS1* (VXXXD) allele (Gill et al., 2018). Likewise, in *Distimake quinquefolius* (Convolvulaceae), the *HSS* locus is polymorphic with alleles encoding VXXXD and VXXXN motifs. A plant homozygous for the VXXXD allele had about twice the concentration of PAs in its tissues as a plant with the VXXXN allele, and, *in vitro*, the VXXXD enzyme had about 20 times the rate of homospermidine production of the VXXXN enzyme (Prakashrao et al., 2022). However, in the latter case, one can also see disconnect between HSS function (reduced 20-fold in the VXXXN HSS) and phenotype (PA concentration reduced only 2-fold) which brings into question how much homospermidine (and HSS optimization) is necessary for PA biosynthesis to evolve. Furthermore, among HSS enzymes of Convolvulaceae, the VXXXD HSS of *D. quinquefolius* (PA positive) has only about twice the specific activity of the IXXXN HSS of *Ipomoea alba* (PA-negative, but producing necine bases that are likely precursors of PAs) (Prakashrao et al., 2022), again bringing into question the predictive value of motif for function and phenotype.

Beyond optimization of HSS catalytic function, substitution of a VXXXD motif for an IXXXN motif reduced DHS function, i.e. deoxyhypusination of eIF5A, *in vitro* (Kaltenegger et al., 2013; Prakashrao et al., 2022). Restoration of DHS function in the IXXXN and VXXXN mutants of the wild-type HSS enzyme of *Parsonsia alboflavescens* (VXXXD) (Table 4) confirms the role of the motif in reduction of DHS function and the primacy of the N273D substitution since the IXXXD mutant, like the wild-type VXXXD enzyme, did not have detectable DHS activity (Table 4). Loss of DHS function may be a prerequisite for evolution of high levels of homospermidine production since DHS is an essential enzyme of primary metabolism (Wator et al., 2020) whose activity should be tightly controlled. Abolition of DHS function would free a paralog to evolve elevated expression levels. However, *Ipomoea alba* (Convolvulaceae) expresses *in vivo* an IXXXN HSS with detectable DHS function *in vitro* (Prakashrao et al., 2022), suggesting that some DHS activity by HSS enzymes is tolerated.

Does the N273D substitutional also increase HSS activity? The IXXXN and VXXXN mutant enzymes both produce about half as much homospermidine as both the wild-type VXXXD HSS and the IXXXD mutant (Table 4). However, since this comparison is based on only four (4) individual enzyme preparations, it does not account for variance in enzyme activity introduced by the preparation process. Multiple replicate preparations of each enzyme would need to be tested to estimate an effect size of this substitution, which is likely to be highly context specific, e.g. mutagenesis of *Ipomoea alba* IXXXN HSS to IXXXD only increased homospermidine production by ca. 20% (Prakashrao et al., 2022).

The functional importance of the I269V substitution in the evolution of HSS is bolstered by evidence of positive selection on this site on the reduced dataset topology (Fig. 4a) in our branch-site analysis (Table 2) and in the selection analyses previously conducted on Convolvulaceae HSS (Kaltenegger et al., 2013). In contrast, we see no evidence that the I269V substitution affects catalytic activity since the IXXXD HSS mutant and the VXXXD wild-type enzyme of *Parsonsia alboflavescens* (Table 4) are equivalent in terms of both homospermidine production and absence of DHS function *in vitro*. Models of the DHS and HSS proteins predict that I269V substitution is located at sites of protein-protein interaction between the monomers that assemble to form the active tetramer (Wator et al., 2020; Prakashrao et al., 2022). As with the N273D substitution, this points to a function in reducing interference with DHS activity (via reducing formation of heterotetramers) rather than increasing HSS activity per se (Kaltenegger and Ober, 2015). But again, the fitness cost of interference with DHS is unclear since IXXXN *HSS*-like genes and DHS genes are expressed in highly overlapping patterns (at the organ level) in multiple Convolvulaceae species (Prakashrao et al., 2022). Likewise, IXXXN HSS-like genes have evolved at least twice in Apocynaceae (Fig 4), and, apart from the pseudogene in *Asclepias syriaca*, most of these genomes retain at least one copy with an intact ORF (Appendix S3).

While all 10 sequenced PA-producing Apocynaceae species have an intact HSS encoding a VXXXD motif (Appendix S1, Table S2), many species with the same motif do not have evidence of PAs (Barny et al., 2021, 2022). Of 17 Apocyneae (VXXXD) genera tested (Barny, et al. 2021), PAs have been found in only two, *Anodendron* and *Amphineurion* (Sasaki and Hirata, 1970; Colegate et al., 2016). Within the MOE clade (all VXXXD, Fig. 1), PAs have been identified with high confidence only in Echiteae and with low confidence in *Mandevilla boliviensis* (Mesechiteae) (Barny et al., 2021). PA occurrence among Malouetieae (all VXXXD) is uncertain; there is low to moderate confidence for their presence in species of *Galactophora*, *Holarrhena*, and *Kibatalia* (Barny et al., 2021). Among Nerieae, *Strophanthus* (VXXXD HSS in *S. boivinii*, *HSS* pseudogene in *S. preussii*) has been much investigated for specialized metabolites, but PAs have never been reported (Knittel et al., 2016). Therefore, while we can predict *HSS* genotype (VXXXD) from PA phenotype with some accuracy in Apocynaceae (and across angiosperms), there is much more ambiguity in predicting phenotype from genotype. Setting aside error in phenotypic classification, see extensive discussion in (Barny et al., 2021, 2022), there are many levels at which this disconnect may occur.

Substitutions at the two focal amino acid sites (positions 269 and 273) may be insufficient to predict an HSS that is optimized for PA production. We identified additional candidate functional sites, homologous to functional sites in human DHS, based on evidence of positive selection (Table 2). Position 19 in Apocynaceae DHS/HSS-like enzymes is predicted to be part of the ball-and-chain motif (Liao et al., 1998; Umland et al., 2004; Wator et al., 2020). Presence of this motif is necessary for DHS to function, but this is the least conserved part of DHS and amino acid changes could accumulate at a higher rate without altering the function of the ball-and-chain motif (Umland et al., 2004; Wator et al., 2020). Five of the amino acids of particular interest (positions 75, 246, 269, 273, 275, identified via selection and/or functional assays) are linked to functional areas in human DHS. Positions 246, 269, 273 and 275 are predicted to be near the enzyme surface at the active site tunnel entrance, so binding substrates are likely to encounter these amino acids. Umland, et al. (2004) showed that the human DHS tunnel entrance had a less negative charge than the modelled *Senecio vulgaris* (Asteraceae) HSS, leading to the prediction that certain amino acid substitutions in this area could affect interactions with the eIF5A precursor protein. Prakashrao et al. (2022) confirmed this polarity difference when comparing models of DHS and HSS of some Convolvulaceae species, but not when comparing *Distimake quinquefolius* DHS (IXXXN) with its two HSS variants: HSS1 (VXXXD) and HSS2 (VXXXN). Relative to the DHS, *S. vulgaris* HSS has the same substitutions at the active site tunnel entrance (L246I, I269V, N273D) as some Apocynaceae HSS. Considering that the putative active sites of all intact Apocynaceae HSS-like amino acid sequences are conserved, it theoretically follows that functional changes could result from amino acid substitutions that deflect bulky eIF5A (i.e. DHS-like function), but allow the entrance of much smaller polyamines (spermidine and putrescine; i.e. HSS-like function) (Umland et al., 2004; Ober and Kaltenegger, 2009; Wator et al., 2020; Prakashrao et al., 2022).

It may be hypothesized that a combination of these sites will more effectively identify a functionally optimized HSS and predict PA phenotype. For example, the L246I substitution creates an extended I[X]_22_VXXXD motif, present in HSS of all Echiteae species and *Amphineurion marginatum* (Apocyneae) (high confidence in PA presence) as well as HSS of *Mandevilla boliviensis* (Mesechiteae), *Eucorymbia alba* (Apocyneae), and *Galactophora schomburgkiana* (Malouetieae) (low to moderate confidence in PA presence) (Barny et al., 2021). *Isonema smeathmanii* also has an I[X]_22_VXXXD motif but its PA phenotype is unknown (Barny et al., 2021). On the other hand, the PA-producing *Anodendron affine* (Apocyneae) (Sasaki and Hirata, 1970) has an L[X]_22_VXXXD motif. Thus, considering an extended motif identifies species that should be further investigated for PAs, but does not obviously predict PA phenotype better than the VXXXD motif alone.

Incongruence between *HSS* genotype (VXXXD) and PA phenotype (negative) could result from mutations in promoter regions that silence *HSS*-like genes with intact ORFs, or these ORFs may be transcribed but not translated as observed in some neotropical bats (Noctilionoidea) where intact S-opsin genes were sequenced from multiple species that had no detectable S-opsin proteins and, by inference, had lost the ability to see in the UV spectrum (Sadier et al., 2018). Sadier et al. (2018) found evidence of multiple independent losses of either transcription or translation of S-opsin genes whose ORFs nevertheless remained intact. The same may be true for *HSS*-like genes of PA-free Apocynaceae. Similarly, loss of the PA phenotype can result from silencing of genes downstream of HSS in the PA biosynthetic pathway since pseudogenization or loss of transcription or translation could occur in any gene of the pathway.

The apparent disconnect between genotype and phenotype could be observed if HSS has other functions besides producing PA precursors, e.g. homospermidine may be a precursor to specialized metabolites other than PAs (Bienz et al., 2002; Prakashrao et al., 2022), or products of HSS catalysis other than homospermidine may be biologically relevant (Fig. 2c). Thus, optimization of HSS function could have predated the evolution of PA biosynthesis. A highly functional HSS may be an exaptation for PA evolution as observed in *Bicyclus anynana* (African squinting bush brown butterfly) where multiple duplications produced the *engrailed* (*en*) gene family, expressed in wing eyespots. However, since these duplications occurred 60 my before the evolution of eyespots, the paralogs were retained for other reasons (Banerjee et al., 2020).

The retention of apparently intact *HSS*-like genes in PA-free Apocynaceae species may indicate that these genes have been re-recruited to DHS function. The DHS in *Trypanosoma brucei* (Euglenozoa) is a heterotetramer comprised of a catalytically impaired DHS paralog (IXXXN), which contains the spermidine binding positions in the active site, and a catalytically dead (VXXXR) paralog, which contains the NAD+ binding sites (Afanador et al., 2018). A heterotetramer comprised of these two paralogs is over 1,000-fold more catalytically active than a homotetramer of the IXXXN DHS (Afanador et al., 2018). The possibility that *HSS* orthologs in Apocynaceae function as *DHS* loci can be tested via *in vitro* assays of heterotetramer formation with DHS monomers and comparing DHS activity between homo- and heterotetramers. The IXXXN *HSS*-like genes of Asclepiadeae and Marsdenieae (Figs. 1, 3, 4), predicted to have re-evolved DHS function based on mutagenesis experiments (Table 4), are of particular interest in this scenario.

Finally, the evolution of a highly functional HSS could hypothetically post-date evolution of the PA biosynthetic pathway. DHS is a promiscuous enzyme that has at least two functions, the secondary of which produces homospermidine (Fig. 2a). Trace amounts of homospermidine, presumably produced by DHS, are ubiquitous in plants (Ober et al., 2003b). The recruitment of a promiscuous enzyme into a novel biosynthetic pathway would occur when the latent product becomes relevant, and this theoretically could occur before, simultaneous to, or after gene duplication (Noda-Garcia and Tawfik, 2020). Therefore, DHS could be recruited to a PA biosynthetic pathway and persist in its substrate promiscuity until specialization of HSS, i.e. inability to bind to eIF5A and increased production of homospermidine, was selected (Kreis and Munkert, 2019; Noda-Garcia and Tawfik, 2020). However, this scenario is not relevant to understanding homology of PA biosynthesis across Apocynaceae since the duplication that gave rise to all functional *HSS* genes in PA-producing species occurred in their common ancestor (Figs 3, 4).

### An *HSS*-like gene with a highly functional motif evolved in the MRCA of all PA-producing species, followed by multiple independent losses, but implications for phenotypic evolution are unclear—

An HSS-like enzyme with an IXXXD motif evolved in the APSA clade after the divergence of Wrightieae (Fig. 4). This is not the VXXXD motif reconstructed by Livshultz, et al. (2018), but this IXXXD enzyme is predicted to have similarly high HSS and low DHS activity as a VXXXD enzyme (Table 4). While this is consistent with origin of PA biosynthesis in this ancestor, there is sufficient evidence that HSS function can be decoupled from PA phenotype (see previous section) to preclude confidence in an inferred phenotype for this ancestor. Evolution of this highly functional HSS-like enzyme early in the diversification of the APSA clade was followed by loss of HSS in 5 independent lineages, evidenced by pseudogenization of all *HSS-*like loci (Figs. 1, 4, Appendix S3). Thus, we know that this *HSS*-like gene did not become essential in all species that inherited it and that there was abundant time for it to become pseudogenized. There were also two independent origins of HSS-like enzymes with, potentially, restored DHS function via re-evolution of the IXXXN motif in Asclepiadeae and in one of two Marsdenieae paralogs (Figs. 3, 4, Appendix S6). And two or four origins of enzymes with VXXXD motifs, and, potentially, even less ability to interfere with DHS function than their IXXXD ancestor (Figs. 3, 4).

### Patterns of selection on HSS do not support a single origin of PAs, but don’t reject it either—

We hypothesized that selection for PAs in an ancestral species would be detectable as positive selection on the *HSS* gene of that ancestor. The ancestral IXXXD HSS (branch A, Fig. 3, 4, S3, S4) evolved in the MRCA of all PA-producing Apocynaceae species (Fig. 1). This gene is reconstructed as being under relaxed selection as evidenced by near neutral ω (0.8, Table 1) and k=0.01 under the RELAX general descriptive model (Fig. 4). Yet, the D75A substitution that maps to this branch (branch A, Appendices S4 and S5) occurred at one of the amino acid positions identified as experiencing significant positive selection in the full dataset tree topology (Table 2). One resolution to this apparent paradox is the high similarity of DHS and HSS at all levels: sequence, structure, function (Prakashrao et al., 2022) such that selection on one or two amino acid positions (undetectable to branch-wide tests) is enough to produce an HSS sufficiently optimized for PA biosynthesis. Alternatively, *de novo* evolution of a highly functional HSS could occur under completely relaxed selection (Weng et al., 2012; Rey et al., 2019). After gene duplication, the duplication-degeneration-complementation (DDC) model predicts that both paralogs will experience relaxation of selection, allowing potentially adaptive mutations to accumulate in one of the paralogs (Force et al., 1999; Ober and Kaltenegger, 2009; Panchy et al., 2016). The ancestral *HSS*-like (Branch A, ω=0.8, k=0.01, Fig. 4) and *DHS*-like (Branch H, ω=0.1, k<0.2, Fig. 4) paralogs both experienced relaxed selection (Appendices S4, S5; Appendix S1: Tables S10, S11), and potentially functionally important amino acid substitutions (D75A, N273D) occurred in the ancestral IXXXD *HSS*. If interference with DHS function (via formation of heterotetramers and/or interaction with eIF5A) is the primary barrier to recruitment of a proto-HSS to novel expression patterns and a novel biosynthetic pathway, then substitutions that degrade DHS function while maintaining HSS function may be the only “adaptations” required. Synthesis of this reconstructed ancestral IXXXD HSS and *in vitro* assays would answer questions about its function, but not if it was involved in a PA biosynthetic pathway.

### Selection tests weakly favor convergent evolution of optimized HSS—

There is weak evidence from selection tests to support convergent evolution of optimized HSS (and possibly PA biosynthesis) in multiple Apocynaceae lineages. The signal of intensifying selection on the entire *HSS*-like clade (clade A) when compared to the *DHS*-like clade (clade H) (k=1.31, Table 3) and a pattern of selection intensification are consistent with multiple origins of functionally optimized HSS (Appendices S4 and S5). However, formal tests of branches where either a VXXXD motif and/or PAs may have evolved identified only one of ten tested branches (F, the ancestral *HSS*-like gene of Malouetieae, Figs. 3, 4) where evolution of the VXXXD motif was accompanied by statistically significant positive selection after correction for multiple testing (Table 1). Presence of PAs among Malouetieae species is still uncertain, but they have been identified with low to moderate confidence in *Holarrhena pubescens, Kibatalia macrophylla,* and *Galactophora schomburgkiana* (Barny, et al. 2021).

Positive selection on *HSS*, however, may not necessarily signal *de novo* evolution of PA biosynthesis but rather HSS optimization under selection from herbivores for increased PA defense. The latter scenario has been proposed for the pattern of selection on threonine deaminase (TD2) in Solanaceae (Rausher and Huang, 2016). Rausher and Huang (2016) suggest that a 25-30 my span of episodic positive selection on TD2 was a result of fluctuating herbivory: high abundance of herbivores selected for adaptive evolution of TD2 and a decrease in herbivore abundance relaxed selection on TD2. Since PAs do not appear to adversely affect adapted specialist herbivores (Cogni et al., 2012; Wei et al., 2015), the cause of fluctuating selection would not be just herbivore abundance but also herbivore community composition. A shift from PA-adapted specialists to PA-susceptible generalists would favor escalation of PA-defense and vice versa (Ali and Agrawal, 2012).

### Limitations of selection tests: potential for incorrect classification of branches—

The validity of our selection tests depends on the correct identification of branches where HSS function either increased or decreased. We used a single motif as a proxy for HSS function, which did not allow us to examine the contributions of other potentially functionally important sites (Table 2), and we have no knowledge of expression levels. We also used the presence of PAs to infer highly functional *HSS* genes. However, the occurrence of PAs is still under-determined in Apocynaceae (Barny et al., 2021), i.e. some species are likely misidentified as lacking PAs, and there is evidence that PAs may be produced with suboptimal HSS enzymes (Prakashrao et al., 2022). Better understanding of structure-function relationships in HSS and PA distribution in Apocynaceae are necessary for future refinement of these analyses. Furthermore, new models may better capture the evolutionary process by simultaneous reconstruction of selection on genotype and phenotype, rather than on genotype alone. A shortcoming of RELAX is the *a priori* designation of foreground and background branches, as is the assumption that trait changes occur at nodes (i.e. speciation events) rather than elsewhere on a branch (Wertheim et al., 2015; Halabi et al., 2020). TraitRELAX increases analytical flexibility by allowing trait transitions to occur multiple times across a branch and uses k (selection intensity parameter) based on these reconstructions to designate foreground and background branches (Halabi, et al. 2020).

### Future work: reducing uncertainty in the genotype-phenotype map to clarify evolution of PA biosynthesis in Apocynaceae—

Despite a detailed reconstruction of the *DHS/HSS* gene tree, there are clearly gaps in data that must be filled to reconstruct evolution of PA biosynthesis in Apocynaceae. The main barrier is uncertainty in the *HSS* genotype to HSS function to PA phenotype map. There is sufficient evidence from infraspecific polymorphisms in other families that catalytically inferior HSS enzymes can still function in PA biosynthesis (Gill et al., 2018; Prakashrao et al., 2022). Thus, understanding when optimal HSS function evolved cannot provide a satisfactory answer to when PAs evolved. Priority is a better understanding of the distribution of PAs in Apocynaceae to allow more confidence in the prediction of PA phenotype from *HSS* genotype. Expression studies are necessary to identify *HSS* pseudogenes that retain intact ORFs. Then, additional candidate genes of the PA biosynthetic pathway need to be identified, e.g. via co-expression network analysis (Wisecaver et al., 2017). The second enzyme, a member of the copper-containing amine oxidase (CuAO) family, has already been identified in PA-producing *Heliotropium indicum* (Boraginaceae) (Zakaria et al., 2022). The identity of this gene and other candidate genes can be predicted by studying expression in relation to presence of PAs (Donoso et al., 2021). Genome mining (Franke et al., 2019) combined with functional assays can then quickly identify functional orthologs across related species. When additional genes in the PA biosynthetic pathway are identified in Apocynaceae, phenotype can be correlated with a multigenic genotype, and evolution of the entire pathway can be reconstructed (Piatkowski et al., 2020). Assembly of an entire biosynthetic pathway provides much stronger evidence for placing the evolution of a metabolite than evolution of a single enzyme of the pathway.

## CONCLUSIONS

We confirm orthology of genes encoding HSS, essential specialist enzymes of PA biosynthesis, among all the scattered species of PA-producing Apocynaceae (Fig. 1). We reconstruct the HSS-like enzyme of their common ancestor with an IXXXD motif (Fig. 4), predicted to have the same high HSS function and reduced DHS function as the VXXXD motif HSS of extant PA-producing species (Table 4). Yet, there is substantiation uncertainty in the mapping of *HSS* genotype to HSS function to PA phenotype when all evidence is synthesized. This precludes firm conclusions about homology of PAs across Apocynaceae and renders ambiguous inferences toward phenotypic evolution from patterns of selection on *HSS*-like genes. We propose that distinguishing between homology of biosynthetic pathways and parallel recruitment of orthologous genes into independently evolved pathways requires detailed understanding of the evolution of the entire pathway, not just one of the component genes.

## ACKNOWLEDGEMENTS

The authors thank Mark Fishbein (Oklahoma State University), André Simões (Universidade Estadual de Campinas), David J. Middleton (Singapore Botanical Garden), and Mary Endress (University of Zurich) for sharing plant leaf tissue. We thank Hobart and William Smith Colleges undergraduates Madison Cullinan, Meredith Steinfeldt, Abbey Foote, Katarina Kostović, and Charmaine Chung for laboratory assistance. This work was supported by the National Science Foundation, United States, grant DEB-1655663 and DEB-1655223 to TL and SCKS and a Drexel University Clinical and Translational Research Institute (CTRI) seed grant to TL.

## DATA AVAILABILITY STATEMENT

The data underlying this article are available in the article, its online supplementary material, and a data supplement archived on Dryad (doi:10.5061/dryad.1c59zw3z2). All sequences are deposited in the GenBank Nucleotide Database (https://www.ncbi.nlm.nih.gov/genbank/) and can be accessed with the accession numbers listed in Appendix S1, Table S2. Probe sequences were published in Straub et al. (2020): the first 2707 sequences in the file here: https://figshare.com/articles/dataset/Apocynaceae_all_bait_sequences_txt/12830579/1.

## ONLINE SUPPORTING INFORMATION

Additional supporting information may be found online in the Supporting Information section at the end of the article.

---

**Appendix S1**. Supplementary tables S1—S13.

**Appendix S2.** Figure S1: Maximum likelihood tree from initial alignment.

**Appendix S3.** Fig. S2: Maximum likelihood tree from full dataset alignment.

**Appendix S4.** Fig. S3: Maximum likelihood tree, reduced dataset topology.

**Appendix S5.** Fig. S4: Maximum likelihood tree, full dataset topology.

**Appendix S6.** Fig. S5: Identification of recombinants among *HSS*-like sequences from Marsdenieae.

**Appendix S7.** Fig. S6: **a**. Confirmation via SDS-PAGE gel eletrophoresis of *Parsonsia alboflavescens* HSS enzyme (wild-type and mutants) expression and purification. **b.** Confirmation via size exclusion chromatography coupled to UV that the enzymes form homotetramers.

**Appendix S8.** Fig. S7: Kinetic curves of *Parsonsia alboflavescens* HSS wild-type and mutant enzymes in the DHS assays versus DHS assay minus eIF5a. Product increase of 1,3-Diaminopropane and Canavalmine is shown.

**Appendix S9.** Fig. S8: Full dataset topology tree with node numbers referenced in the PAML reconstruction of ancestral sequences and in aBSREL and RELAX estimations of ω and k in Appendix S1: Tables S7 and S10.

**Appendix S10.** Fig. S9: Reduced dataset tree with node numbers referenced in the PAML reconstruction of ancestral sequences and in aBSREL and RELAX estimations of ω and k in Appendix S1: Tables S8 and S11.

**Appendix S2 Figure S1.**
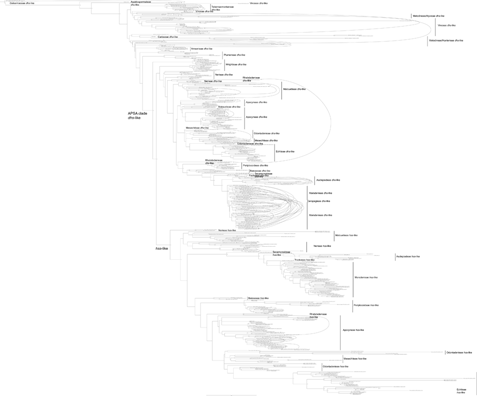
Maximum likelihood tree from initial alignment (see ^85^ Methods: Gene assembly from contigs; Appendix S1: Table S3), including all assembled contigs and previously sequenced DHS-like and HSS-like genes. Arrows link contigs concatenated for downstream analyses. Contigs with non-terminal stop codons marked by “Ψ” in the sequence name. Bootstrap support values >50% mapped at nodes.

**Appendix S3 Fig. S2:**
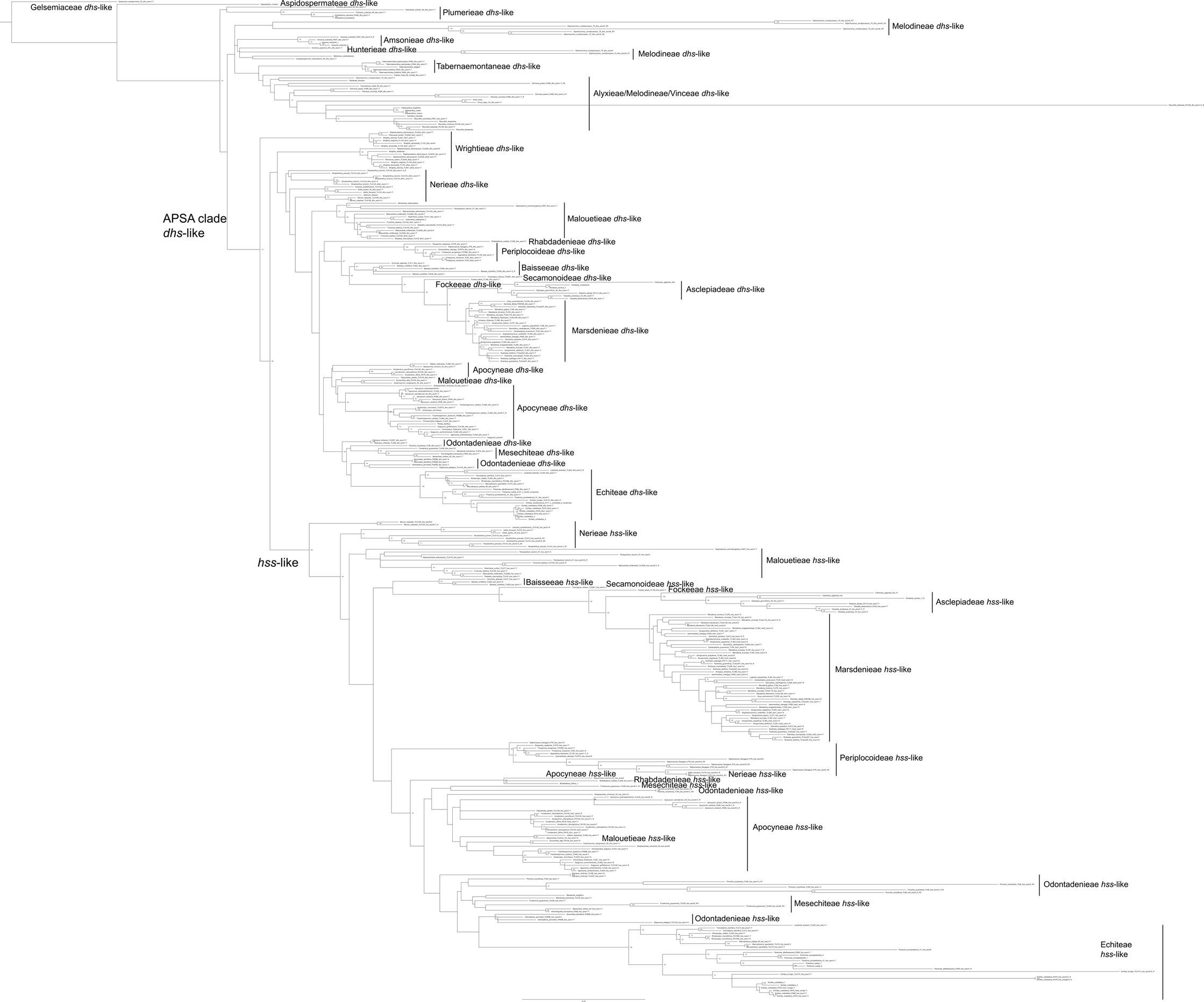
Maximum likelihood tree from full dataset alignment (see Methods: Gene assembly from contigs, Construction of matrices, Appendix S1: Table S3), including all fully assembled gene sequences, concatenated contigs, additional short contigs, potential pseudogene contigs, and previously sequenced DHS-like and HSS-like genes. Contigs with non-terminal stop codons marked by “Ψ” in the sequence name. Bootstrap support values >50% mapped at nodes.

**Appendix S4 Fig. S3:**
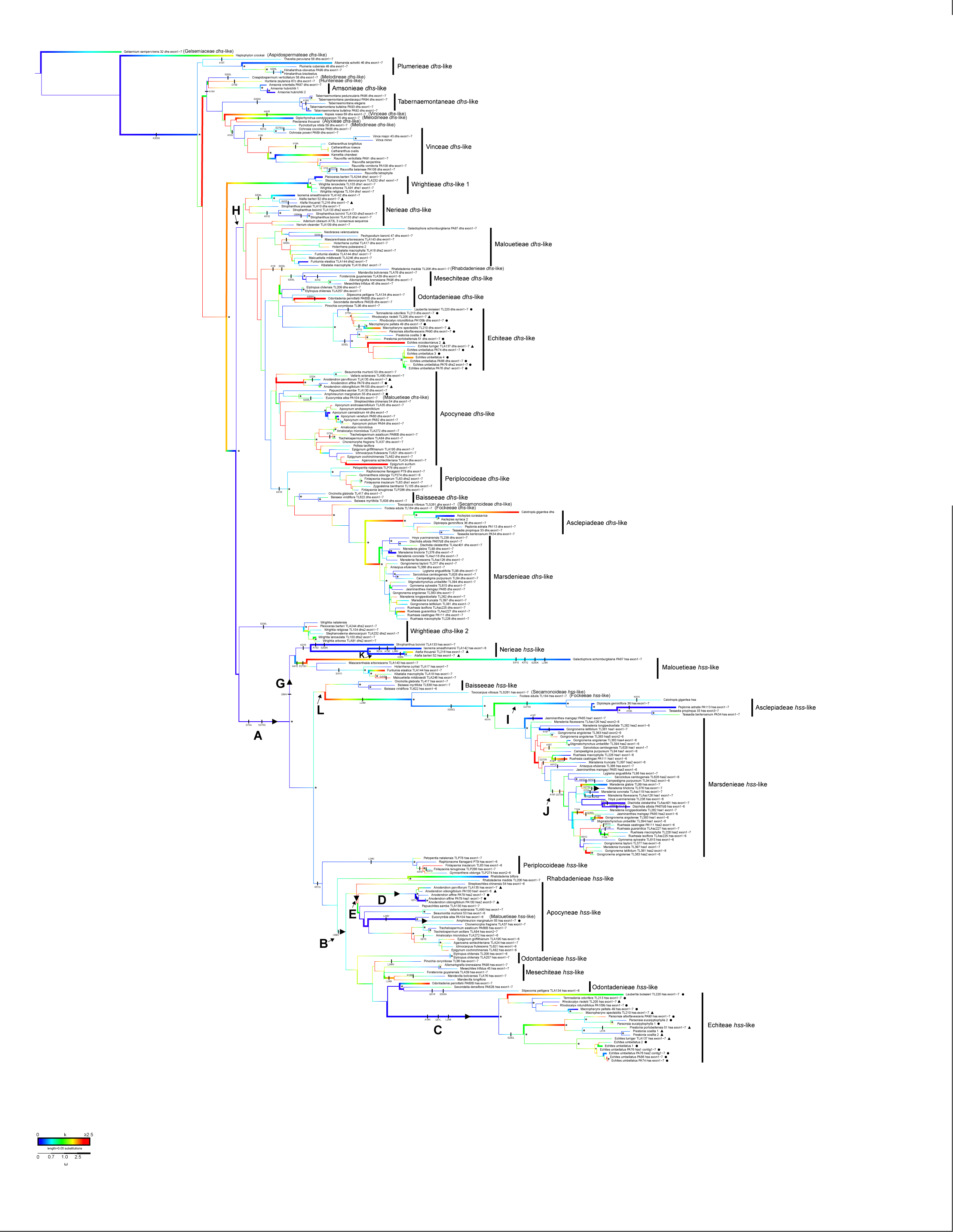
Maximum likelihood tree, reduced dataset topology (see Methods: Gene Tree Construction, Figs. 3a, c, 4a for summary trees). Branch lengths are proportional to probability of nucleotide substitution. Branch widths are proportional to the relative rate of nonsynonymous to synonymous amino acid changes (ω), divided into four categories: 1) ω<0.7, purifying selection; 2) 0.7< ω < 1.0, neutral evolution; 3) 1< ω < 2.5, slightly positive selection; 4) ω >2.5, strongly positive selection. Branch color depicts the selection intensity parameter, k, calculated under the RELAX descriptive model (see Methods: Tests for selection), with darkest blue corresponding to k=0, i.e. ω1, ω2, ω3= 1, complete relaxation of selection, and darkest red to k>=2.5, i.e. ω1<<1, ω2=1, ω3>>1, intensification of selection. Substitutions at amino acid positions experiencing episodic diversifying selection (see Table 2) and in the I269VXXXN273D motif are mapped on the branches. Branches and lineages A-L, used in selection tests, (see Fig. 3, Tables 1-3) are labeled with letters and arrowheads. Presence of PAs is indicated next to all sequences from that species (i.e. both DHS- and HSS-like sequences): circles correspond to PA presence in the species with high confidence, as defined by Barny et al. (2021); triangles correspond to PA presence in the genus with high confidence. Bootstrap support > 90 % is indicated with asterisks.

**Appendix S5 Fig. S4:**
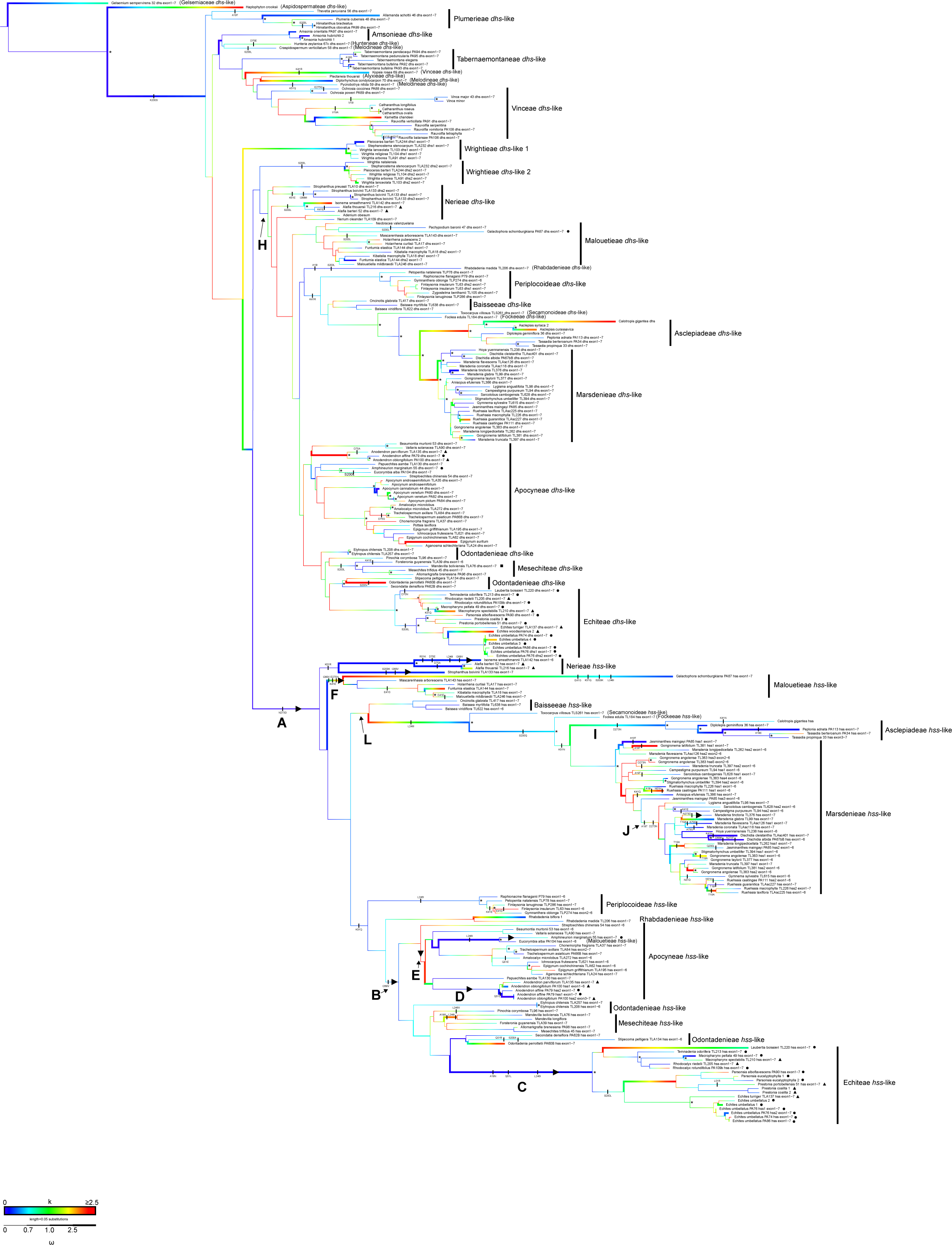
Maximum likelihood tree, full dataset topology [see Methods: Gene Tree Construction: reduced dataset alignment, constrained to the tree topology from the full dataset alignment (Appendix S3: Fig. S2); Figs. 3b, d, 4b for summary trees]. Branch lengths are proportional to probability of nucleotide substitution. Branch widths are proportional to the relative rate of nonsynonymous to synonymous amino acid changes (ω), divided into four categories: 1) ω<0.7, purifying selection; 2) 0.7< ω < 1.0, neutral evolution; 3) 1< ω < 2.5, slightly positive selection; 4) ω >2.5, strongly positive selection. Branch color depicts the selection intensity parameter, k, calculated under the RELAX descriptive model (see Methods: Tests for selection), with darkest blue corresponding to k=0, i.e. ω1, ω2, ω3= 1, complete relaxation of selection, and darkest red to k>=2.5, i.e. ω1<<1, ω2=1, ω3>>1, intensification of selection. Substitutions at amino acid positions experiencing episodic diversifying selection (see Table 2) and in the I269VXXXN273D motif are mapped on the branches. Branches and lineages A-L, used in selection tests, (see Fig. 3, Tables 1-3) are labeled with letters and arrowheads. Presence of PAs is indicated next to all sequences from that species (i.e. both DHS- and HSS-like sequences): circles correspond to PA presence in the species with high confidence, as defined by Barny et al. (2021); triangles correspond to PA presence in the genus with high confidence. Bootstrap support > 90 % is indicated with asterisks.

**Appendix S6 Fig. S5:**
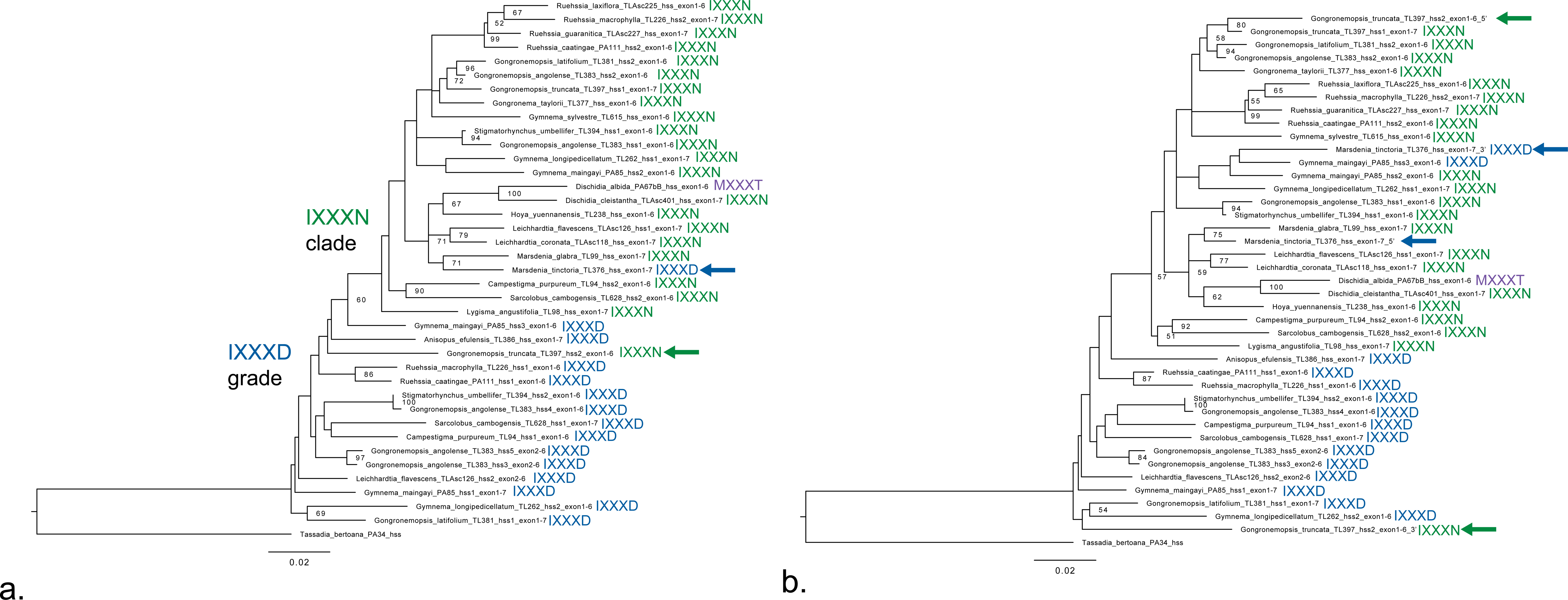
Identification of recombinants among HSS-like sequences from Marsdenieae. a. ML topology (See Methods: Gene tree construction and Tests for recombination) of the full Marsdenieae HSS-like sequences showing anomalous placement of the *Mardenia tinctoria* sequence (IXXXD motif) in the IXXXN clade and the Gongronemopsis truncata sequence (IXXXN motif) among IXXXD sequences. b. ML topology with the 5’ and 3’ ends of the M. tinctoria and G. truncata sequences included as distinct terminals, separated at a likely recombination breakpoint identified with GARD (See Methods: Tests for recombination). The *Gongronemopsis truncata* sequence is likely a recombinant between a sequence in the IXXXN clade and the IXXXD grade, but the anomalous position of the 3’ end with the IXXXN motif is not caused by recombination. The *Marsdenia tinctoria* sequence is not a recombinant. Bootstrap values >50 displayed at nodes and amino acid motif at positions 269 and 273 are noted next to sequence names.

**Appendix S7. Fig. S6:**
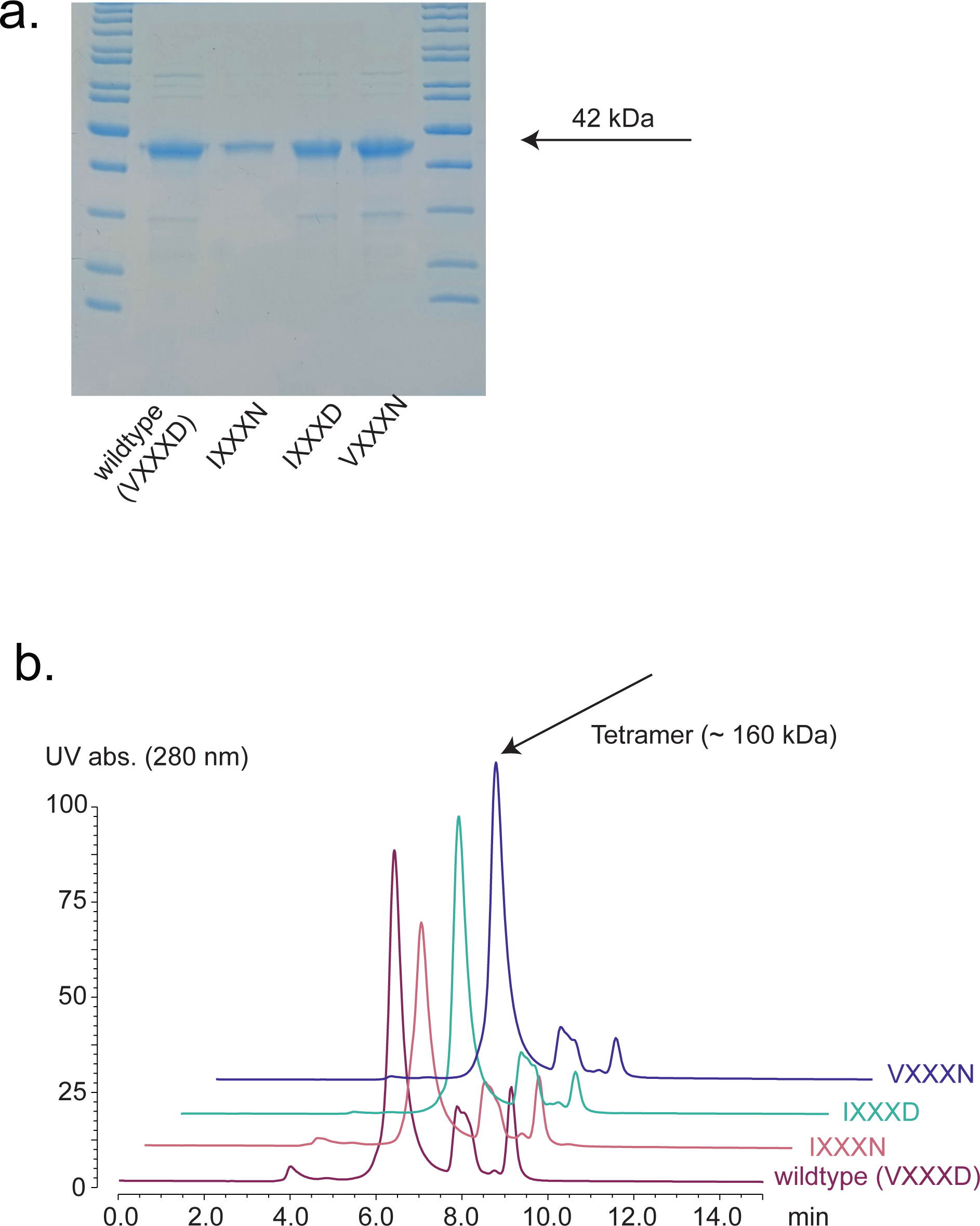
a. Confirmation via SDS-PAGE gel eletrophoresis of Parsonsia alboflavescens HSS enzyme (wild-type and mutants) expression and purification. b. Confirmation via size exclusion chromatography coupled to UV that the enzymes form homotetramers.

**Appendix S8 Fig. S7:**
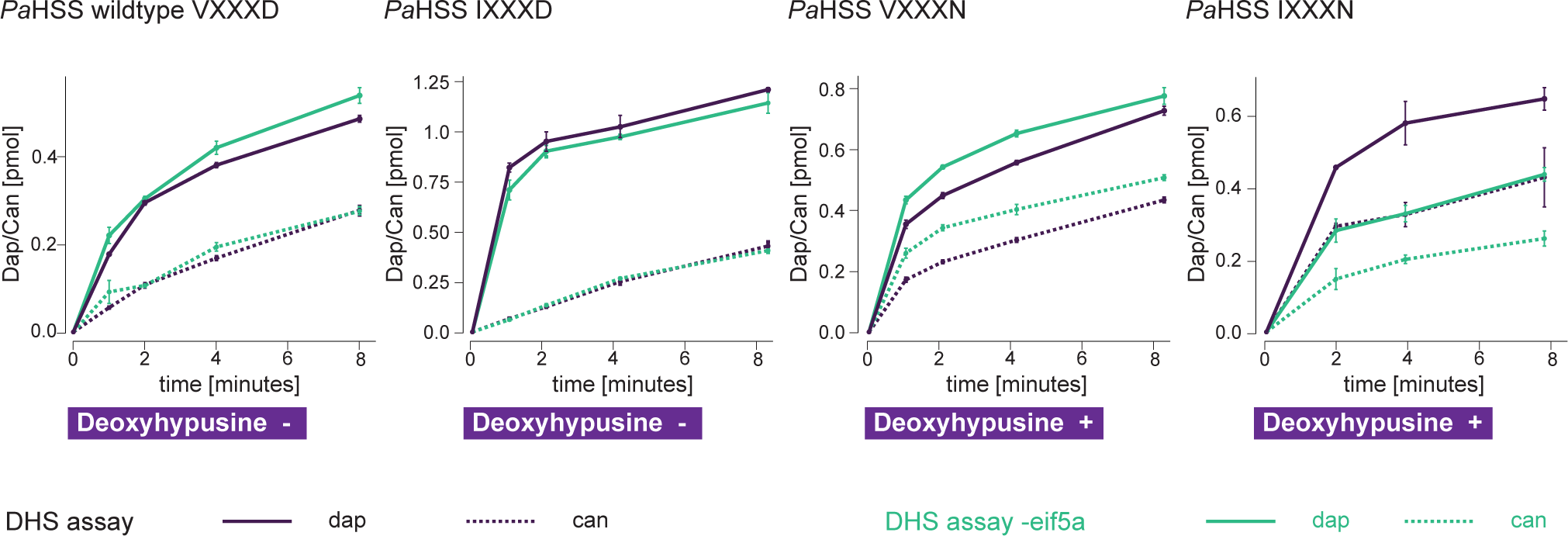
Kinetic curves of Parsonsia alboflavescens HSS wildtype and mutant enzymes in the DHS assays versus DHS assay minus eIF5a. Product increase of 1,3-Diaminopropane and Canavalmine is shown.

**Appendix S9 Fig. S8:**
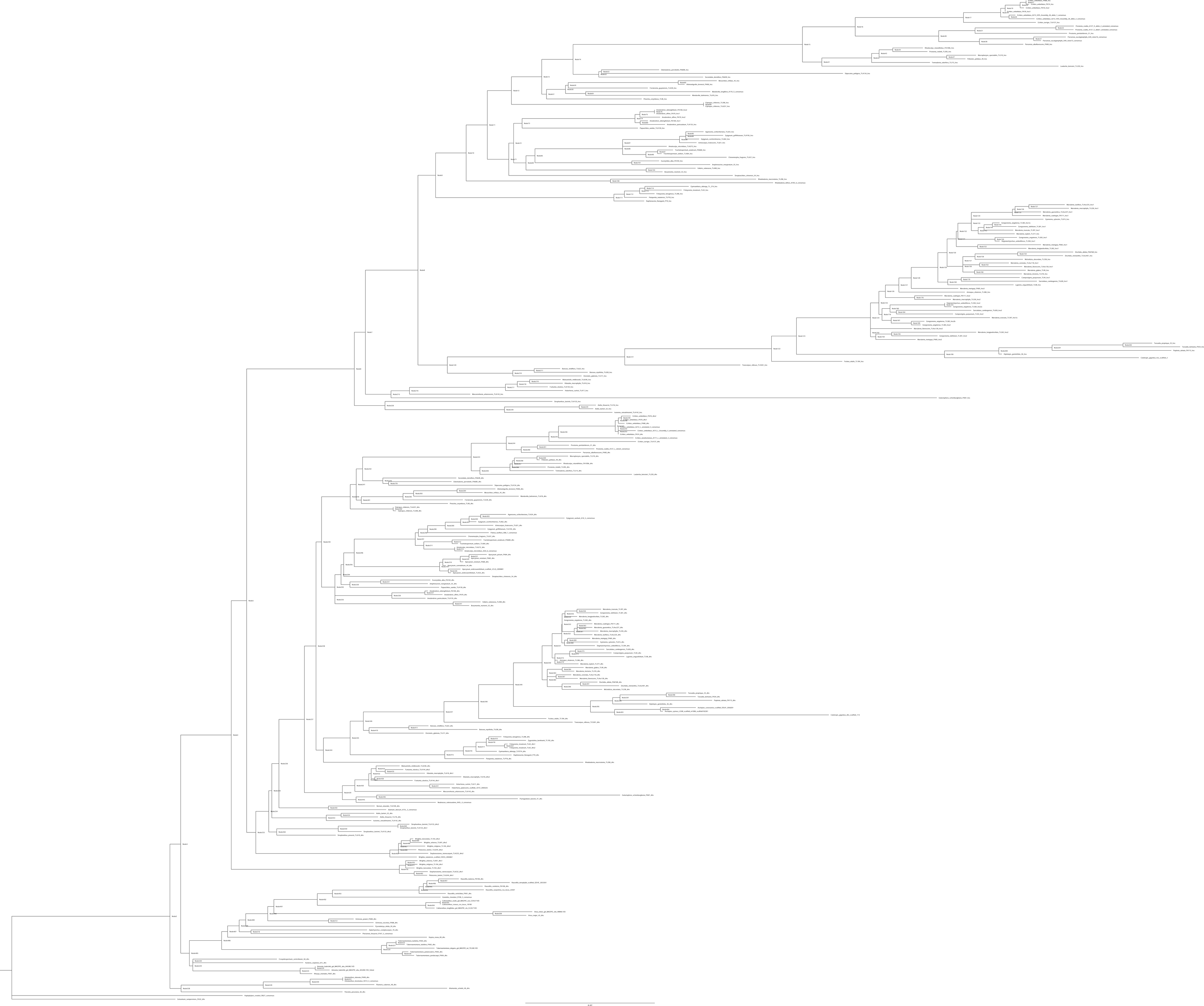
Full dataset topology tree with node numbers referenced in the PAML reconstruction of ancestral sequences and in aBSREL and RELAX estimations of ωand k in Appendix S1: Tables S7 and S10. Smith et al.— American Journal of Botany 2024 – Appendix S10

**Appendix S10 Fig. S9:**
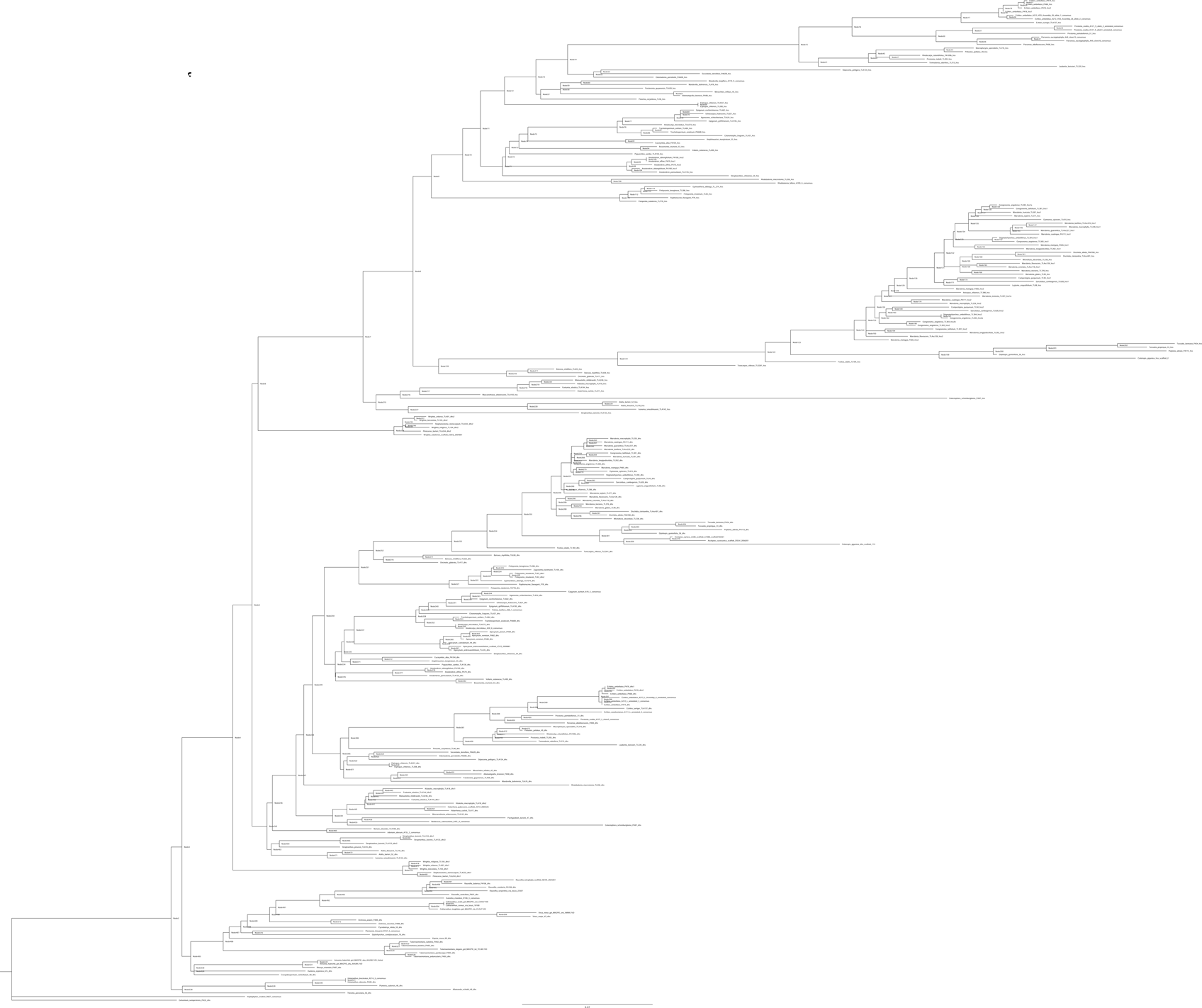
Reduced dataset tree with node numbers referenced in the PAML reconstruction of ancestral sequences and in aBSREL and RELAX estimations of ω and k in Appendix S1: Tables S8 and S11.

